# O-GlcNAc modification forces the formation of an α-Synuclein amyloid-strain with notably diminished seeding activity and pathology

**DOI:** 10.1101/2023.03.07.531573

**Authors:** Aaron T. Balana, Anne-Laure Mahul-Mellier, Binh A Nguyen, Mian Horvath, Afraah Javed, Eldon R. Hard, Yllza Jasiqi, Preeti Singh, Shumaila Afrin, Rose Pedretti, Virender Singh, Virginia M.-Y. Lee, Kelvin C. Luk, Lorena Saelices, Hilal A. Lashuel, Matthew R. Pratt

## Abstract

The process of amyloid fibril formation remains one of the primary targets for developing diagnostics and treatments for several neurodegenerative diseases (NDDs). Amyloid-forming proteins such α-Synuclein and Tau, which are implicated in the pathogenesis of Alzheimer’s and Parkinson’s disease, can form different types of fibril structure, or strains, that exhibit distinct structures, toxic properties, seeding activities, and pathology spreading patterns in the brain. Therefore, understanding the molecular and structural determinants contributing to the formation of different amyloid strains or their distinct features could open new avenues for developing disease-specific diagnostics and therapies. In this work, we report that O-GlcNAc modification of α-Synuclein monomers results in the formation of amyloid fibril with distinct core structure, as revealed by Cryo-EM, and diminished seeding activity in seeding-based neuronal and rodent models of Parkinson’s disease. Although the mechanisms underpinning the seeding neutralization activity of the O-GlcNAc modified fibrils remain unclear, our *in vitro* mechanistic studies indicate that heat shock proteins interactions with O-GlcNAc fibril inhibit their seeding activity, suggesting that the O-GlcNAc modification may alter the interactome of the α-Synuclein fibrils in ways that lead to reduce seeding activity in vivo. Our results show that post-translational modifications, such as O-GlcNAc modification, of α-Synuclein are key determinants of α-Synuclein amyloid strains and pathogenicity. These findings have significant implications for how we investigate and target amyloids in the brain and could possibly explain the lack of correlation between amyloid burden and neurodegeneration or cognitive decline in some subtypes of NDDs.

## INTRODUCTION

The formation and deposition of misfolded protein aggregates in the brain is a common feature of most neurodegenerative diseases (NDDs)^1,2^. Peptides and proteins that are entirely or partially unfolded in solution are particularly prone to misfolding and formation of β-sheet rich fibrillar aggregates characterized by the eventual stacking of monomers into β-sheets in a cross-β conformation, also known as amyloid fibrils^3^. The formation of these aggregates is associated with cell dysfunction and death. Several forms of misfolded protein oligomers and mature fibrils of different morphology and structural properties are toxic to mammalian cells and neurons in culture^4^. Additionally, amyloid fibrils derived from several proteins implicated in the pathogenesis of NDDs, such as Tau, α-Synuclein (α-Syn), and TDP43, efficiently seed and induce the aggregation of the corresponding endogenous protein when added to neurons^5–7^ and spread through cell-to-cell propagation mechanisms to different brain regions when injected into the brain of rodents^8–12^. These observations, combined with the strong genetic and neuropathological evidence linking the aggregation of these proteins to the pathogenesis of NDDs, continue to drive strong interest in targeting the formation, pathogenic properties, or removal of these amyloid structures as potential therapeutic strategies for the treatment of NDDs, as evidence by the increasing number of anti-aggregation therapies in different phases of clinical trials today.

The ability of these individual proteins to form multiple amyloid structures or “strains” in different NDDs presents significant challenges to current anti-amyloid therapies and amyloid-based diagnostics, as they have not accounted for the structural and biochemical diversity of the different strains^13–15^. For example, α-Syn amyloid fibrils isolated from Parkinson’s disease (PD), multiple system atrophy (MSA), or Lewy body dementia (LBD) patients exhibit distinct structures^16–18^. Injection of α-Syn aggregates isolated from patients with different diseases results in distinct phenotypes and pathology patterns in mice^19^. These results have generated significant interest in uncovering structure-pathogenicity relationships between these strains and the molecular and structural determinants of their formation. Addressing this knowledge gap could lead to new strategies to develop disease-specific diagnostics (e.g., PET-tracers) and therapies or the identification of generalized strategies to neutralize the pathogenic activities of all α-Syn fibril strains.

Posttranslational modifications (PTMs) can have profound consequences on the structure, biochemistry, and function of proteins in health and disease. Biochemical studies of pathological hallmarks of NDDs, including amyloid plaques, tangles, Lewy bodies (LB), and Lewy neurites, have shown that they accumulate misfolded Aβ, Tau, and α-Syn aggregates that are subjected to different types of PTMs at multiple residues^20–24^. Consequently, antibodies against the modified and aggregate forms of these proteins have become the main tools to detect, monitor, and quantify pathology formation in post-mortem human brain tissues and animal models of NDDs. Most PTMs are substoichiometric in nature and may not be found in high percentages on the aggregates from patients. However, the prion-like nature of amyloids has potential to give them outsized influence. These findings, combined with increasing evidence demonstrating an increase in the levels of specific modified forms of α-Syn or Tau in patients with PD and Alzheimer’s disease (AD), respectively, suggest that PTMs may play an important role in regulating pathology formation and neurodegeneration in NDDs. Despite this, several questions regarding the roles of PTMs in NDDs remain unanswered, including 1) which PTMs enhance or protect against protein aggregation and toxicity in NDDs; 2) how do PTMs influence the structural, biochemical, and cellular properties of fibrils; and 3) do they play critical roles in regulating α-Syn seeding and pathology spreading?

In this work, we sought to address this knowledge gap by conducting systematic studies to determine how one specific PTM, an intracellular form of glycosylation called O-GlcNAc (Figure 1a)^25^, influences the structural properties and seeding activity of α-Syn in vitro, in neurons, and animal models of α-Syn pathology formation and spreading. O-GlcNAc modifications are the addition of monosaccharide *N*-acetylglucosamine to the side chains of serine and threonine residues. O-GlcNAc can dynamically cycle on and off proteins through the action of two enzymes: O-GlcNAc transferase (OGT) that adds the modifications and O-GlcNAcase (OGA) that removes them. O-GlcNAc has been phenotypically linked to several biological processes, including protein aggregation and neurodegeneration in several NDDs^26,27^. For example, O-GlcNAc levels are 40-50% lower in AD brains when compared to age-matched controls^28^, and neuron-specific knockout of OGT in mice results in tau hyperphosphorylation and neurodegeneration^29^. Notably, both tau and α-Syn are O-GlcNAc modified at multiple positions *in vivo* (Figure 1b).

**Figure 1.**
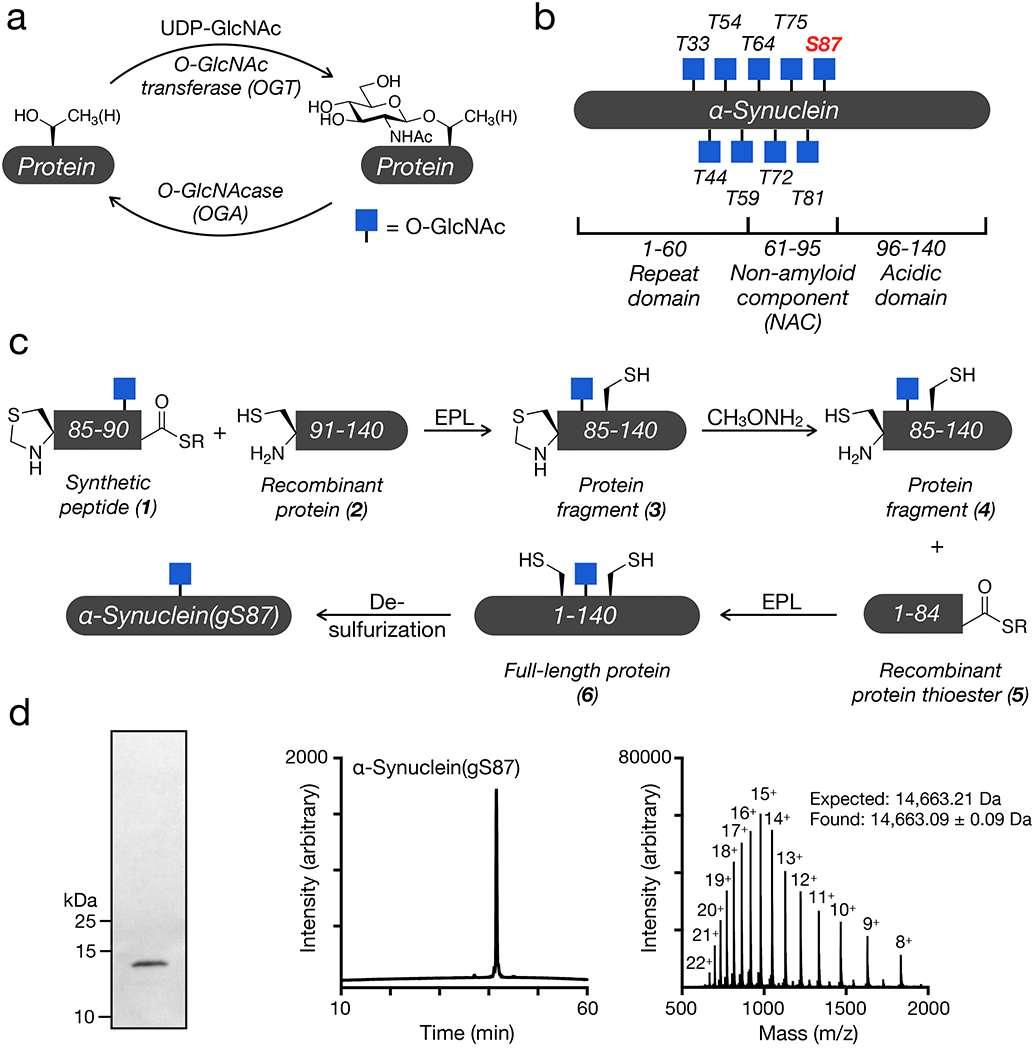
O-GlcNAc modified α-Syn. a) O-GlcNAc is the dynamic addition of N-acetylglucosamine to serine and threonine residues of intracellular proteins. b) α-Syn is O-GlcNAc modified *in vivo* at several residues that can alter its amyloid aggregation. The focus of this study, O-GlcNAc on serine 87 (gS87), is highlighted in red. c) Synthesis of α-Syn(gS87) using expressed protein ligation. d) Characterization of α-Syn(gS87) by gel electrophoresis, high-performance liquid chromatography (HPLC), and mass spectrometry.

These and other observations have led to the creation of a range of OGA inhibitors that can elevate O-GlcNAc modifications as potential therapeutics^30^. Multiple pre-clinical studies in animal models of AD and PD demonstrated that OGA inhibition increases O-GlcNAc in brains and slows the formation of amyloid aggregates and neuron death^31,32^. Additionally, a recent analysis of α-Syn in brain tissue from the Line 61 mouse-model that overexpresses human α-Syn brain found that 20% of the protein is O-GlcNAc modified and that these levels rise to ∼35% upon OGA inhibition^32^. Some of these compounds have advanced to the clinic and show no overt toxicity in humans. We and others have shown that O-GlcNAc on tau and α-Syn can directly slow the kinetics of amyloid aggregation of these proteins *in vitro* in a site-specific fashion^31,33–35^. These data indicate that O-GlcNAc may protect neurons by inhibiting amyloid aggregation and that increasing the levels of this modification with drugs may slow the progression of certain neurodegenerative diseases.

In our previous studies on α-Syn O-GlcNAc modification^34,35^, we used synthetic protein chemistry to prepare the protein bearing O-GlcNAc at four different sites (T72, T75, T81 or S87). Specifically, we took advantage of expressed protein ligation (EPL)^36^, an extension of native chemical ligation (NCL), as these methods are the only way to install O-GlcNAc in a site- and stereo-specific fashion^37^. Subsequent biochemical analysis showed that individual O-GlcNAc residues slow the kinetics of α-Syn aggregation with notable site-specific differences. However, these O-GlcNAc modifications did not completely stop the formation of α-Syn aggregates over time. This led us to hypothesize that O-GlcNAc may alter the progression of PD by not only slowing the rate of amyloid formation but by causing the formation of amyloid strains with altered pathogenicity and toxicity.

Here, we applied protein synthesis in combination with biochemical, cellular, *in vivo*, and structural analyses to test this hypothesis. We focused on O-GlcNAc modification at S87, termed α-Syn(gS87), because this modification-site displays the least inhibition of α-Syn fibril formation, thus providing an opportunity to obtain homogeneously O-GlcNAc-modified fibrils for detailed analysis. First, we used protein semisynthesis to generate large amounts of α-Syn(gS87). We then used a variety of *in vitro* experiments to show that α-Syn(gS87) does form amyloid fibrils that have a different core structure and that these fibrils seed the aggregation of monomeric α-Syn and template their structure onto the newly-formed wild-type (WT) α-Syn aggregates. Most importantly, we observed that α-Syn(gS87) fibrils failed to induce toxicity or α-Syn aggregation in primary neurons (as detected by phosphorylation of α-Syn at S129). These results were recapitulated *in vivo* using the well-established seeding-mediate mouse models of α-Syn aggregation and pathology spreading^7,12^. Systematic mechanistic studies on α-Syn seeding in neurons in both α-Syn knock-out (KO) and WT primary neurons demonstrated that this lack of seeding activity and pathology formation do not result from differential uptake, degradation, or processing of α-Syn(gS87) amyloids compared to unmodified protein. Rather, we observe an interesting divergence in the behavior of the α-Syn(gS87) fibrils, where they can seed aggregation *in vitro* but not in neurons or mice. We demonstrate that this may be due to differential recognition of the fiber strains by heat shock proteins, as HSP27 is better able to inhibit aggregation seeded by α-Syn(gS87) fibrils *in vitro*. Finally, we used cryo-EM to obtain a structural model of the α-Syn(gS87) amyloid that is very different from both other in vitro fibrils and ex vivo aggregates from patients.

Taken together, our results confirm that O-GlcNAcylation can force the formation of a novel α-Syn amyloid strain with diminished pathogenicity in neurons and *in vivo*. This adds even more evidence for a model where O-GlcNAc may not only slow the aggregation of α-Syn, but could also protect against the progression of neurodegenerative diseases through multiple mechanisms. To our knowledge, this α-Syn strain is also the first amyloid that is capable of strongly-seeding aggregation *in vitro* but does not in neurons. It also appears to be the first example of a PTM that can almost completely block the seeding and spread of α-Syn *in vivo*. More broadly, our results also demonstrate that some amyloids may be almost benign despite their ability to seed aggregation *in vitro*, with important implications for amyloid characterization and highlighting the importance of therapeutically-targeting the correct and pathogenic structure in associated diseases.

## RESULTS

### Synthesis of α-Syn(gS87)

As stated above, we used an EPL-based strategy to prepare α-Syn(gS87) (Figure 1c). EPL involves the chemoselective reaction, termed native chemical ligation (NCL), between a protein-thioester and another protein fragment bearing an N-terminal cysteine residue to form a native amide-bond. Accordingly, we first used solid-phase peptide synthesis to prepare peptide thioester **1** (residues 85-90) containing an O-GlcNAc at serine 87. We then recombinantly expressed protein **2** (residues 91-140) and performed an EPL reaction with **1** to yield α-Syn(gS87) fragment **3**. The corresponding N-terminal thiazolidine of **3** was then removed to unmask the N-terminal cysteine in **4**. After recombinant production of protein thioester **5** (residues 1-84) using an intein fusion, we performed another EPL reaction to yield full-length α-Syn(gS87) **6**. α-Syn contains no native cysteine residues that can be exploited for EPL. However, we could simply perform desulfurization(Wan and Danishefsky, 2007) of the cysteine residues in **6** to transform them into the alanines found at residues 85 and 91 in α-Syn and yielding α-Syn(gS87) with no primary sequences mutations. We then used a combination of techniques to confirm the purity and identity of our synthetic protein (Figure 1d).

### α-Syn(gS87) forms amyloid aggregates with a qualitatively different overall structure

We previously demonstrated that monomeric α-Syn(gS87) forms fibrillar aggregates, although with reduced kinetics compared to the unmodified protein^35^. In those studies, we used a low concentration (50 μM) of α-Syn(gS87) monomers, but others have shown that higher concentrations of α-Syn more efficiently facilitate the production of fibrils, hereafter referred to as pre-formed fibrils (PFFs), in large amounts suitable for subsequent *in vitro*, neuron culture, and *in vivo* seeding experiments. Therefore, we separately subjected α-Syn or α-Syn(gS87) to the well-established and validated α-Syn fibrillization conditions, i.e., incubation of α-Syn (50 or 172 μM) for 7 days in phosphate buffered silane (PBS)^38^. The extent of fibril formation was monitored by Thioflavin T (ThT) fluorescence, and the final fibril preparations were characterized by transmission electron microscopy (TEM), sedimentation assays^39^, and limited proteolysis with proteinase K (PK) (Figure 2a). As expected, α-Syn(gS87) resulted in indistinguishable levels of ThT signal at both concentrations. Notably, others have found that ThT-binding alone is not sufficient to allow for direct comparison of the extent of fibril formation because amyloid fibrils can bind ThT differently, yielding higher or lower signals^40,41^. To account for this possibility, we also analyzed the aggregation reactions by sedimentation, SDS-PAGE, and Coomassie staining (Figure 2b). These data confirmed that α-Syn(gS87) aggregates but to a lesser extent, consistent with our previous findings demonstrating that O-GlcNAc inhibits the nucleation step of α-Syn aggregation^35^.

**Figure 2.**
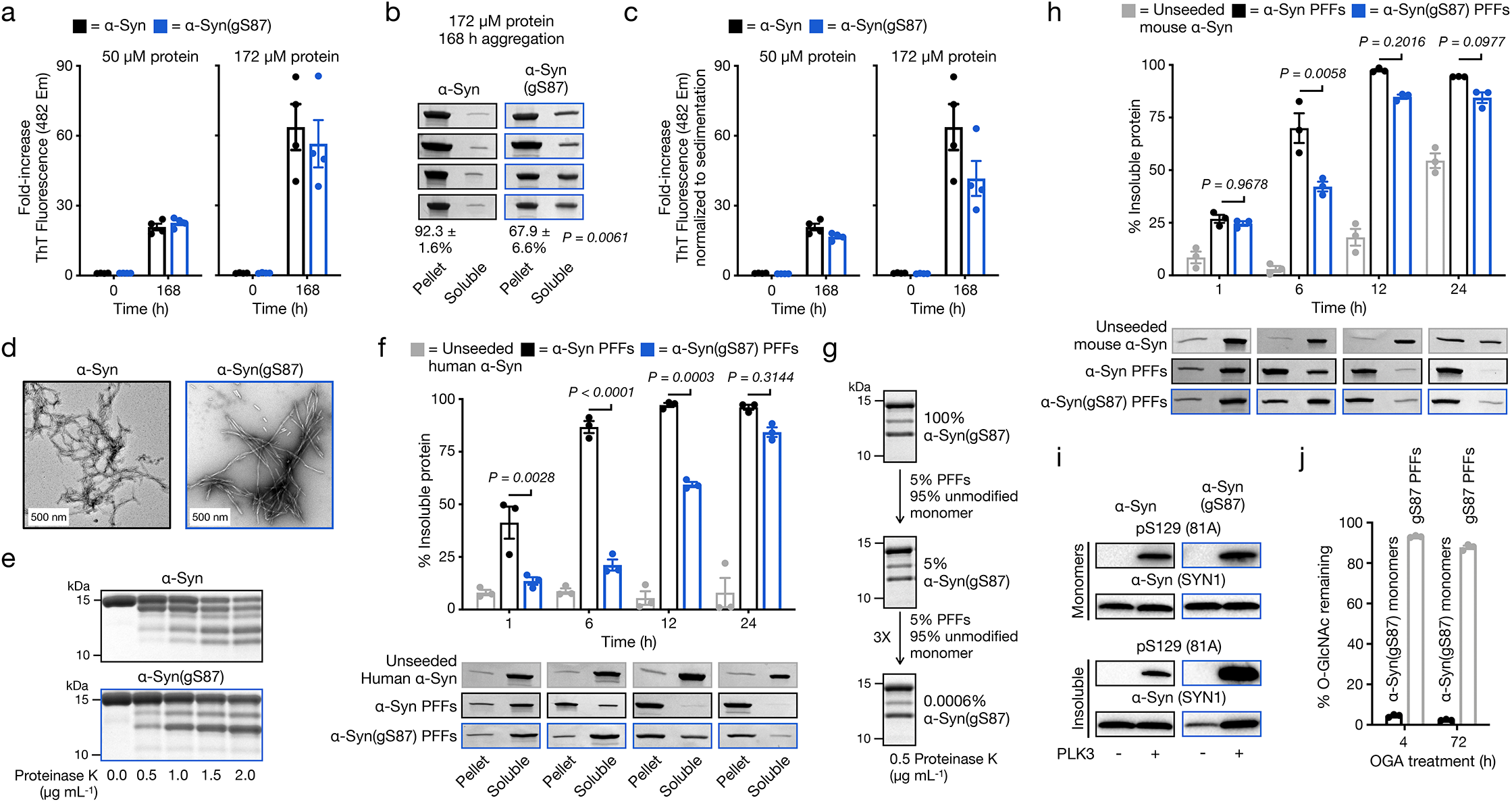
In vitro generation and characterization of α-Syn(gS87) fibrils. a) α-Syn(gS87) aggregates into ThT-positive amyloids. α-Syn and α-Syn(gS87) were subjected to aggregation conditions at the indicated concentrations before analysis by ThT fluorescence. The y-axis shows a fold change in fluorescence compared with α-Syn alone at t = 0 h. Results are mean ±SEM of experimental replicates (n=4). b) α-Syn(gS87) aggregates can be quantitated by sedimentation. α-Syn and α-Syn(gS87) were subjected to aggregation conditions followed by sedimentation and visualization by Coomassie staining. Results are mean ±SEM of experimental replicates (n=4). c) ThT data from (a) normalized to the sedimentation levels in (b). d) The aggregation reactions in (a) (172 μM) were analyzed by TEM after 168 h. e) α-Syn(gS87) forms fibrillar aggregates of distinct structures as discerned from proteinase-K (PK) digestion. The aggregation reactions in (a) (172 μM) were subjected to the indicated concentrations of PK for 30 min before separation by SDS-PAGE and visualization by Coomassie staining. f) α-Syn(gS87) PFFs can seed aggregation of unmodified, human α-Syn. α-Syn PFFs (unmodified or gS87) were added to buffer or unmodified, human α-Syn (50 μM monomer concentration, 5% PFF) before aggregation and analysis by sedimentation and Coomassie staining. Results are mean ±SEM of experimental replicates (n=3). g) α-Syn(gS87) amyloids template their structure onto unmodified, human WT α-Syn. α-Syn was iteratively seeded and the amyloid structure analyzed by PK digestion. h) α-Syn(gS87) PFFs can seed the aggregation of unmodified, mouse α-Syn. α-Syn PFFs (unmodified or gS87) were added to buffer or unmodified, mouse α-Syn (50 μM monomer concentration, 5% PFF) before aggregation and analysis by sedimentation and Coomassie staining. Results are mean ±SEM of experimental replicates (n=3). i) α-Syn(gS87) can be phosphorylated at serine 129 (pS129). The indicated α-Syn PFFs or monomers were incubated with PLK3, and pS129 on α-Syn was visualized by western blotting (WB) using pS129 antibody (81a). j) The O-GlcNAc on α-Syn(gS87) is resistant to removal by OGA post-aggregation. α-Syn(gS87) monomers or PFFs were treated with OGA for the indicated amounts of time, and the removal of O-GlcNAc was measured using RP-HPLC. Results are mean ±SEM of experimental replicates (n=3).

Normalization of the ThT signal using this sedimentation data (Figure 2c) indicated that α-Syn(gS87) could be forming an amyloid strain with slightly higher ThT binding compared to α-Syn. We then examined the aggregates using TEM, confirming that both α-Syn and α-Syn(gS87) are forming amyloid fibrils (Figure 2d). To indirectly probe for structural differences between the two types of fibrils, we subjected unmodified and α-Syn(gS87) to proteinase K (PK) digestion and compared their PK-resistant core sequences by western blotting (WB). Briefly, PK readily digests monomeric α-Syn to very small peptide fragments; however, it cannot gain access to the core of amyloid fibrils resulting in limited proteolysis. We visualized the PK digestion reactions by SDS-PAGE and Coomassie staining and found the expected 5 bands associated with the typical PFF structure of α-Syn, while observing only 3 bands from α-Syn(gS87) PFFs. Together, these results show that α-Syn(gS87) can indeed form an amyloid strain that differs from those formed by the unmodified protein.

### α-Syn(gS87) seeds aggregation and is suitable for cellular and *in vivo* analyses

Next, we asked whether α-Syn(gS87) PFFs could seed the aggregation of unmodified protein. We first generated either α-Syn or α-Syn(gS87) PFFs by incubation of the monomeric forms of these proteins for 7 days (172 μM). Next, we compared the ability of the two PFF preparations to seed the aggregation of human, unmodified α-Syn (50 μM with 5% PFFs) *in vitro*. The efficiency of seeding was assessed by monitoring the kinetics of aggregation by sedimentation as above and found that α-Syn(gS87) PFFs do seed monomeric protein (Figure 2f & Figure S1). Although fibril formation is slower in the presence of α-Syn(gS87) PFFs compared to α-Syn PFFs, the extent of fibrillization is indistinguishable by 24 h. We subjected a portion of the α-Syn(gS87) seeded aggregation reaction to PK digestion and found that α-Syn(gS87) PFFs template their amyloid structure onto unmodified protein (Figure 2g). Furthermore, we discovered that this templating behavior is maintained over multiple rounds of seeding, resulting in the same 3-band PK digestion pattern despite the presence of only 0.0006% remaining α-Syn(gS87) in the amyloids (Figure 2g).

PFF treatment of mouse neurons or injection of α-Syn PFFs into mice brains represent standard approaches to evaluate PFF pathogenicity, toxicity, and propagation^38^. However, previous studies have shown that human α-Syn PFFs seed mouse α-Syn monomers less efficiently than mouse α-Syn PFFs^7,12,42^. This raised the possibility that our human α-Syn(gS87) PFFs may not seed the aggregation of mouse α-Syn. Therefore, we tested whether α-Syn(gS87) PFFs can seed mouse α-Syn aggregation in vitro and found that they can and reach similar levels after only 12 h (Figure 2h & Figure S2). We and others have also shown that the seeding of endogenous α-Syn aggregation in neurons results in phosphorylation of the newly seeded aggregates at serine 129 (pS129)^7^, as observed in the human PD pathology. This enables pS129 to be used as a surrogate marker for pathology formation and propagation in neurons and *in vivo*. We reasoned that the α-Syn(gS87) aggregate structure might be refractory to phosphorylation. Since this structure is templated (Figure 2g), it might prevent us from visualizing neuronal seeding by α-Syn(gS87) PFFs using a pS129 antibody. To determine if α-Syn(gS87) can be phosphorylated at pS129, we incubated monomers and PFFs with PLK3, a kinase that contributes to pS129. We found that α-Syn(gS87) PFFs can be phosphorylated, indicating that we can still use pS129 as a mark for any pathology induced by these PFFs. Finally, because O-GlcNAc is a dynamic modification, we were concerned that cellular OGA might remove O-GlcNAc once the PFFs were taken up into neurons, complicating our analysis. As expected, when we treated α-Syn(gS87) monomers with OGA in vitro, we observed rapid loss of the O-GlcNAc modification (Figure 2j & Figure S3). However, O-GlcNAc on the α-Syn(gS87) PFFs was quite stable, even after 72 h of OGA treatment (Figure 2j & Figure S3), suggesting that it is inaccessible and potentially buried in the amyloid structure. Overall, these data show that α-Syn(gS87) PFFs seed and template their structure onto monomeric, unmodified protein and can be analyzed using standard methods in mouse neurons and brains.

### α-Syn(gS87) PFFs induce significantly less seeding and pathology in neurons and *in vivo*

Next, we investigated the effect of O-GlcNAc modification at S87 on α-Syn-PFFs-mediated induction α-Syn fibrillization and formation of LB-like inclusions using the well established seeding-dependent neuronal model of α-Syn pathology formation^5^. Towards this goal, we compared the seeding activity and seeding-dependent neuron loss of α-Syn(gS87) PFFs and α-Syn PFFs. Specifically, hippocampal primary neurons from mouse embryos were cultured for 7 days in vitro (DIV) and then treated with different concentrations of either α-Syn or α-Syn(gS87) PFFs for 12 days. We then measured the extent of PFFs-mediated seeding of α-Syn aggregation and neuron viability by quantitation of the corresponding fluorophore-labeled antibodies against pS129 and NeuN, respectively (Figure 3a). Consistent with previous data, α-Syn PFFs induced robust seeding activity and aggregation of endogenous α-Syn, and neuron death. In stark contrast, in neurons treated with α-Syn(gS87) PFFs, we did not observe any increase in pS129 signal or neuron death compared to neurons treated with PBS (negative control).

**Figure 3.**
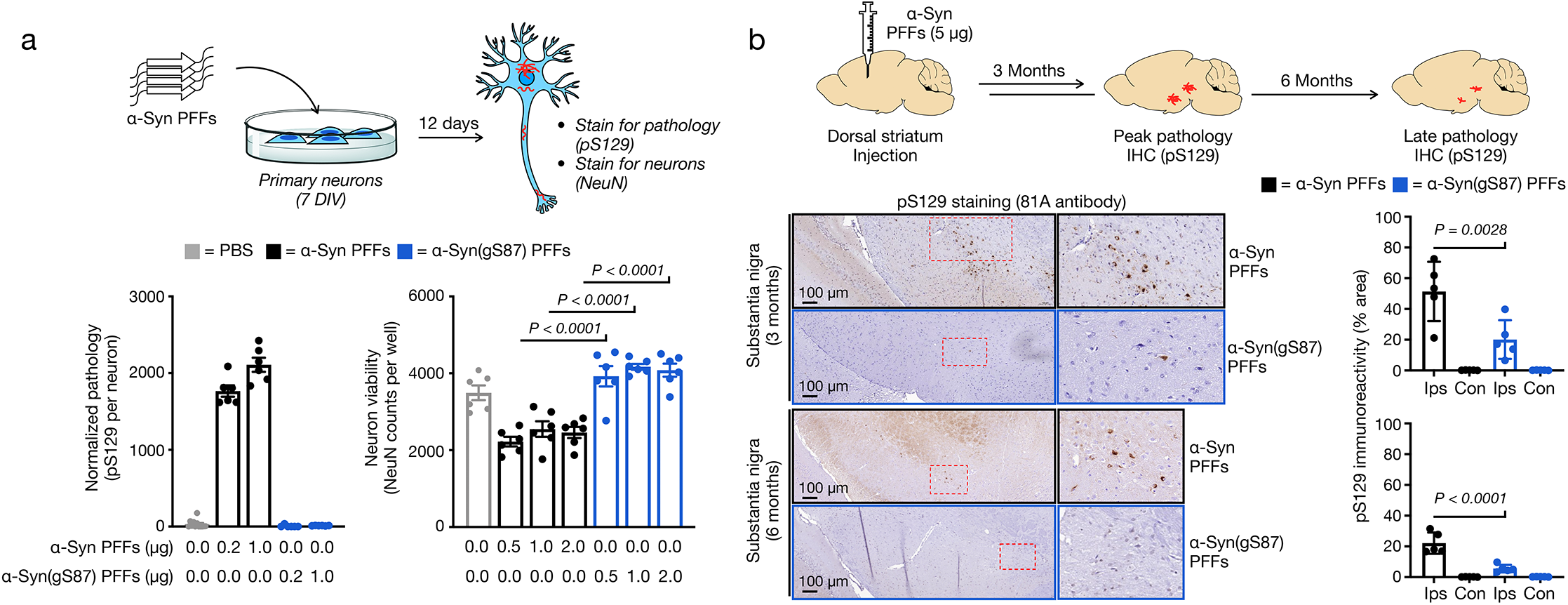
α-Syn(gS87) PFFs have diminished ability to induce pathology formation, spreading, and toxicity in cells and in vivo. a) Primary embryonic hippocampal neurons at 7 days in vitro (7 DIV) were treated with the indicated concentrations of PFFs or PBS for 12 days before analysis of pathology (pS129 staining) and neuron viability (NeuN staining). Results are mean ±SEM of biological replicates (n=6). Statistical significance was determined using a one-way ANOVA test followed by Sidak’s multiple comparison test. b) Wild-type mice were injected with α-Syn or α-Syn(gS87) PFFs (5 μg) in a single unilateral injection into the dorsal striatum. Pathology was visualized by immunohistochemistry against pS129 at 3 and 6 months post-injection. Results are mean ±SEM of biological replicates (n=5). Statistical significance was determined using a one-way ANOVA test followed by Tukey’s multiple comparison test.

Given the striking nature of this difference, we sought to validate these findings in vivo. Towards this goal, we performed a dorsal striatum injection of either α-Syn or α-Syn(gS87) PFFs (5 μg) into the brains of wild-type C57Bl6/C3H mice. After 3 and 6 months, we sacrificed the mice and stained for pS129 as a marker for α-Syn aggregation and Lewy body pathology. In this model, injection of human α-Syn PFFs results in peak pathology at around 3 months that diminishes over time and no significant loss of tyrosine-hydroxylase (TH) positive neurons in the Substantia Nigra^12^. Consistent with our results from cultured neurons, we observed dramatically less overall pS129 staining in the Substantia Nigra and significantly less area with pS129-positive inclusions (Figure 3b). We found the same difference in other parts of the brain including the amygdala and motor cortex (Figure S4). We reasoned that one possible explanation for these differences could be the complete loss of neurons in the α-Syn(gS87) PFF-injected mice. We ruled this possibility out, however, by observing no significant loss of TH-positive neurons in mice injected with either α-Syn or α-Syn(gS87) PFFs (Figure S4). These data demonstrate that α-Syn(gS87) PFFs are less prone to induce pS129 pathology in neurons and *in vivo*. They also set up an interesting divergence between our in vitro data where α-Syn(gS87) PFFs can seed additional aggregation (Figure 2f & h) and in neurons where they exhibit dramatically reduced seeding activity.

### Unmodified and α-Syn(gS87) PFFs have similar uptake, processing, and stability in neurons

The spread of α-Syn PFFs and induction of further aggregation and toxicity is a multistep process^7,43^. In culture, PFFs are first taken up by neurons through the endosomal/lysosomal pathway. After they gain access to the cytosol, the PFFs are cleaved by calpain (residues 114 and 122 *in vitro*)^6,7,44^ and potentially other proteases to generate truncated amyloids with a monomeric molecular weight of ∼12 kDa, down from the 15 kDa of full-length α-Syn. These truncated PFFs seed the aggregation of endogenous α-Syn, which is then phosphorylated to give the pS129 mark associated with pathology. Finally, the aggregates mature into LB-like structures made up of proteins, mostly α-Syn, p62 and ubiquitin, as well as lipids, membranous structures and organelles. This overall pathway is associated with neuron dysfunction and cell death. Neuronal dysfunction coincides with α-Syn fibrillization, but neuron death is more prominent during the transition from fibrils to LB-like inclusions^7,39^. We reasoned that a better understanding of α-Syn(gS87) PFFs structural properties, internalization, processing and seeding in neurons could provide mechanistic insight that could explain why they can seed and template aggregation *in vitro* but apparently not in neurons or brains.

First, in-depth characterization of the unmodified α-Syn and α-Syn(gS87) PFFs revealed that they are quite similar in that they are both made up of primarily fibrils over oligomers and are of a similar length distribution suitable for uptake by neurons (Figure S5). To determine if the two types of fibrils are differentially taken up by neurons, we quantified their uptake in primary neurons from α-Syn KO mice^6,7^, as they allow us to specifically investigate the fate of exogenous unmodified and α-Syn(gS87) PFFs, without confounding issues due to the presence or aggregation of the endogenous protein (Figure 4a).

**Figure 4.**
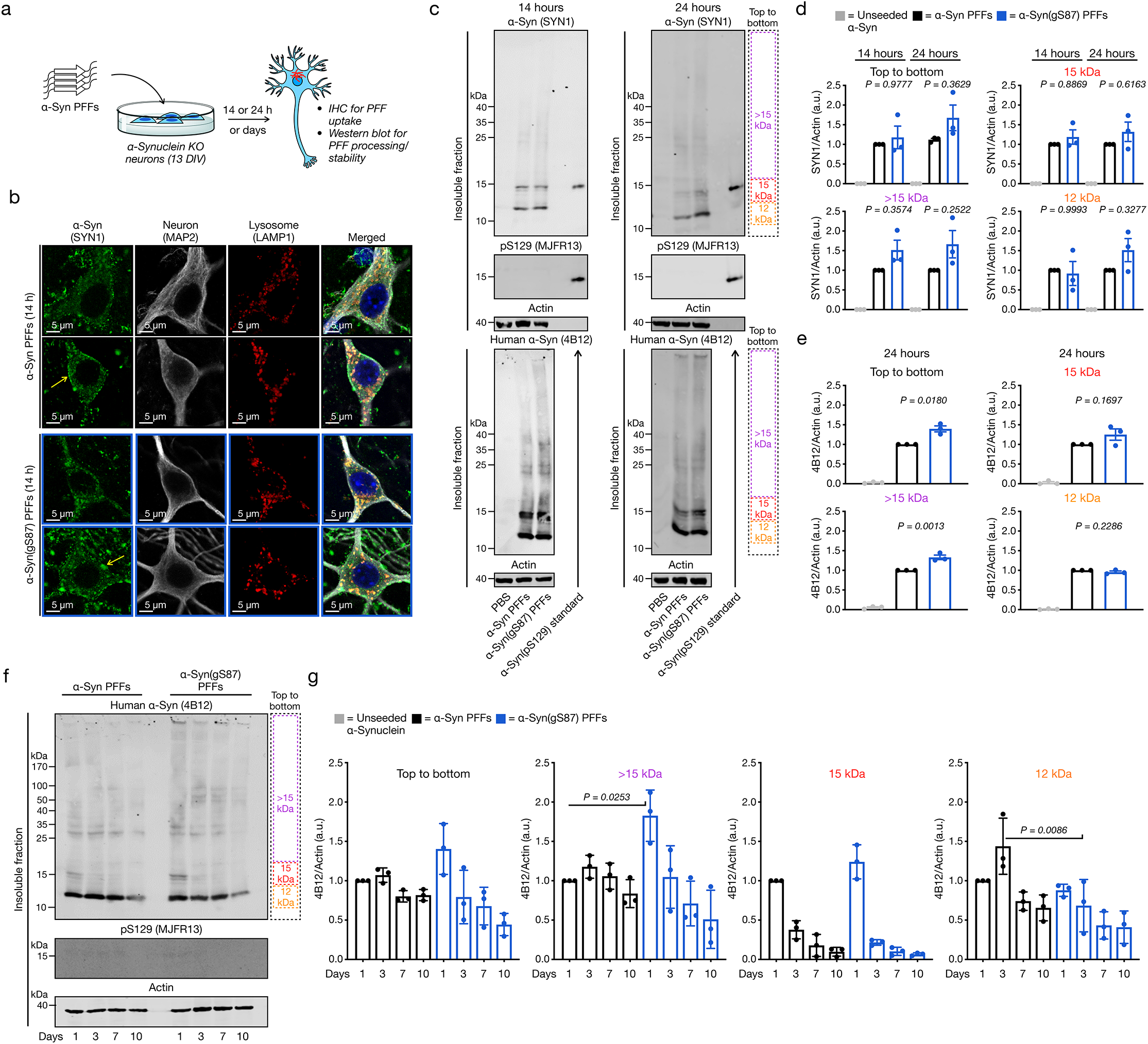
Unmodified and α-Syn(gS87) PFFs display similar uptake, processing, and stability in neurons. a) Experimental outline. Primary hippocampal neurons from α-Syn KO mice at 13 days in vitro (13 DIV) were treated with the indicated PFFs (70 nM) or PBS for different lengths of time before the following analyses. b) O-GlcNAc at S87 does not alter the internalization of PFFs as visualized by immunocytochemistry (ICC) after 14 h of treatment. c) O-GlcNAc at S87 ldoes not significantly affect significantly the internalization, C-terminal cleavage to ∼12 kDa fragment, or phosphorylation at S129 (pS129) of PFFs as visualized by WB after 14 or 24 h of treatment. d) Quantitation of the data in (c) using the pan-α-Syn antibody SYN1. Results are mean ±SEM of biological replicates (n=3). e) Quantitation of the data in (c) using the human-specific α-Syn antibody 4B12. Results are mean ±SEM of biological replicates (n=3). f) O-GlcNAc at S87 does not notably alter the stability of internalized PFFs as visualized by WB over 10 days of treatment. g) Quantitation of the data in (f). Results are mean ±SEM of biological replicates (n=3). In all experiments, statistical significance was determined using a one-way ANOVA test followed by Tukey’s multiple comparison test.

We assessed the uptake of PFFs by treating these neurons with α-Syn or α-Syn(gS87) PFFs (70 nM) for 14 h and examined their localization using immunocytochemistry (ICC) (Figure 4b). As expected^6,45^, we found that unmodified PFFs and α-Syn(gS87) PFFs accumulated on cellular membranes (yellow arrows) and were taken up into neurons and appeared as puncta structure (Figure 4b). Both PFFs seeds strongly colocalized with the lysosomes (LAMP1 positive-vesicles) (Figure 4b), indicating that the O-GlcNAc modification does not influence the internalization of the α-Syn PFFs via the endosomal/lysosomal pathway. Next, we investigated whether α-Syn(gS87) PFFs were processed differently by treating α-Syn KO neurons for 14 or 24 h. WB analysis of the insoluble fraction of the PFF-treated neurons was performed using both pan (SYN1) and human-specific (4B12) α-Syn antibodies. Figure 4c confirmed the internalization of the PFFs in the neurons as indicated by the presence of SDS-resistant aggregates above 15 kDa (HMWs, > 15 kDa) to a similar level (Figure 4d-e) in the unmodified or α-Syn(gS87) PFFs-treated α-Syn KO neurons. As previously described^6,7^, the unmodified PFFs were efficiently cleaved at the C-terminus during the 24 hours post-treatment leading to the accumulation of a 12 kDa fragment, and these seeds were not phosphorylated at S129 (pS129) following their C-terminal cleavage (Figure 4c). Notably, we observed no differences in the proteolytic processing or phosphorylation of the α-Syn(gS87) PFFs (Figure 4c). To confirm these observations, we performed multiple biological replicates (Figure S6) and quantified the WB signal from different regions of the blot (Figure 4d). Using this analysis, we found no significant differences between α-Syn(gS87) PFFs using the pan-antibody (SYN1) and only small differences with the human-specific antibody (4B12). Mouse PFFs were used as a positive control in all of our experiments, and no striking differences in terms of internalization and processing were observed compared to the unmodified and α-Syn(gS87) PFFs (Figure S6). Finally, we sought to explore if the two types of PFFs were differentially cleared once internalized into neurons. Unmodified and α-Syn(gS87) PFFs (70 nM) were added to α-Syn KO neurons, and their fate was monitored for up to 10 days by measuring the amount of remaining PFFs by WB (Figure 4f). Interestingly, we observed a trend where the internalized α-Syn(gS87) PFFs appear to be cleared faster than the unmodified α-Syn(gS87) PFFs, which we again confirmed by performing multiple repeat experiments (Figure S7) (Figure 4g). Together, these results demonstrate that O-GlcNAc modification of α-Syn does not alter uptake, proteolytic processing, direct phosphorylation or the exogenous α-Syn PFFs, but may promote their clearance in neurons (Figure 4g).

### α-Syn(gS87) PFFs seed far less aggregation in neurons compared to unmodified PFFs

Given that we did not observe any major differences in the uptake, processing, or cellular stability between the two types of PFFs, we next examined whether O-GlcNac modification of α-Syn fibrils influences their ability to seed the aggregation of endogenous α-Syn or the formation of LB-like inclusions in neurons. Specifically, we treated primary neurons from wild-type mouse pups with α-Syn or α-Syn(gS87) PFFs (70 nM) for 14 or 21 days and examined the extent of seeding and formation of LB-like inclusions using ICC combined with high-content imaging (HCA) quantification and WB analyses (Figure 5a)^7^. Consistent with our data in Figure 3a, we again observed almost no detectable α-Syn(gS87)-PFF-induced pS129 staining after 14 days and significantly less at 21 days compared to unmodified PFFs. These two sets of results (Figures 3a and 5b), obtained by independent researchers at different locations and using two different model systems (embryonic versus post-natal neurons), confirm that α-Syn(gS87) PFFs exhibit diminished seeding activity and induced much less pathology as measured by pS129. To further validate our findings, we isolated the insoluble and soluble fractions of the neurons and analyzed them by WB (Figure 5c). We observed a ladder of protein species detected by both anti-α-Syn and anti-pS129 antibodies in neurons treated with unmodified PFFs, consistent with the partial stability of the seeded aggregates to SDS. In contrast, we found very little laddering and a much lower overall WB signal from neurons treated with α-Syn(gS87) PFFs. We repeated this experiment two more times (Figure S8), and quantitation of the blots confirmed that α-Syn(gS87) PFFs seed significantly less aggregation of endogenous α-Syn (Figure 5c). We also confirmed that mouse PFFs induce an indistinguishable WB pattern compared to human, unmodified PFFs (Figure S8), as previously described^39^. We then analyzed the portion of α-Syn that remained in the soluble-protein fraction (Figures 5c and S8). As expected, we found that treatment with unmodified α-Syn PFFs results in the loss of soluble α-Syn as it is consumed by the seeded aggregation process. However, in the case of α-Syn(gS87) PFF treatment, we observed essentially no significant loss of soluble α-Syn, which is consistent with their diminished ability to seed endogenous α-Syn in neurons.

**Figure 5.**
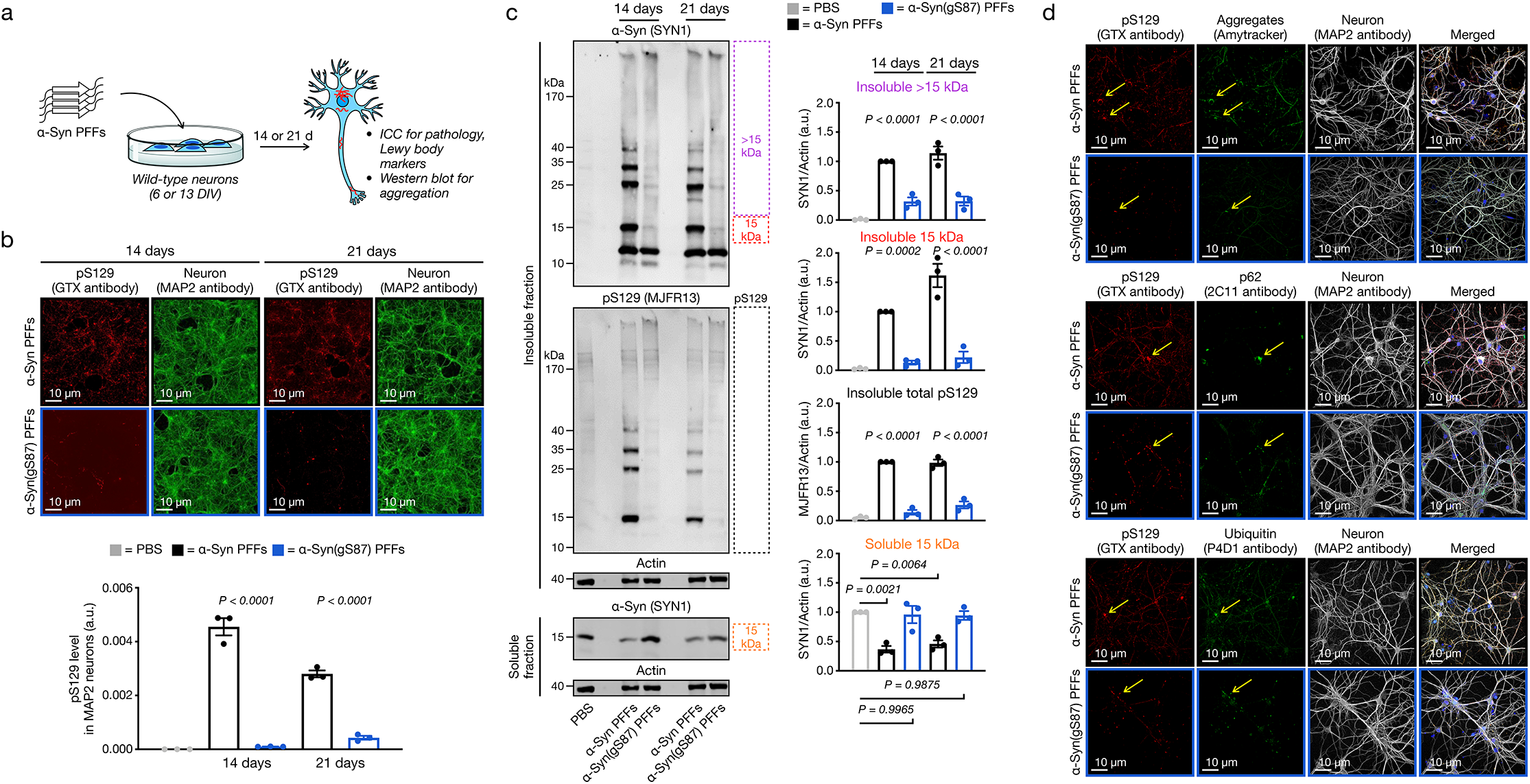
α-Syn(gS87) PFFs have dramatically reduced seeding capacity in neurons. a) Experimental outline. Primary hippocampal neurons from wild-type mice at 6 or 13 days in vitro (DIV 6 or 13) were treated with the indicated PFFs (70 nM) or PBS for 21 and 14 days, respectively, before the following analyses. b) O-GlcNAc at S87 notably reduced the formation of pS129-positive aggregates in neurons as visualized and quantified using immunocytochemistry (ICC) combined to high content imaging. c) Unmodified PFFs seed the aggregation of endogenous α-Syn into insoluble and pS129-positive higher molecular-weight aggregates. O-GlcNAc at S87 dramatically reduced this seeded aggregation, and more endogenous α-Syn remained soluble. Results shown in b and c are the mean ±SD of biological replicates (n=3). Statistical significance was determined using a one-way ANOVA test followed by Tukey’s multiple comparison test. d) Aggregates that form from α-Syn(gS87) PFFs are notably reduced but display Lewy body hallmarks (amyloid, p62 & ubiquitination) by immunocytochemistry (ICC).

We then investigated by ICC and confocal imaging, whether the seeded-aggregates formed in α-Syn(gS87) PFFs-treated neurons were differently stained by the well-established LB markers, including the autophagy-marker p62, ubiquitin (ub) and the Amytracker amyloid-like specific dye (Figures 5d and S9). First, all the pS129-positive seeded aggregates formed in the α-Syn(gS87) PFFs-treated neurons were positively stained by p62, ub and the Amytracker dye, indicating that when these aggregates do form, they share the same hallmarks of the seeded aggregates formed in the unmodified PFFs-treated neurons. Secondly, no LB-like inclusions were observed in α-Syn(gS87) PFFs-treated neurons after 21 days of treatment. We previously reported that this type of inclusions is usually not observed in neurons treated with human unmodified PFF^39^. This observation suggests that α-Syn(gS87) PFFs do not unlock their ability to induce LB-like inclusion formation, as was recently observed for the human α-Syn E83Q PFFs^39^. Finally, we observed that when the aggregates were formed in the α-Syn(gS87) PFFs-treated neurons, they mostly localized in the soma. Barely any aggregates were observed in the neurites of these neurons compared to the extensive neuritic pathology observed in unmodified PFF-treated neurons (Figures 5d).

Overall, these data show that although α-Syn(gS87) PFFs can seed aggregation of unmodified α-Syn *in vitro*, they exhibit diminished seeding activity in living neurons. Additionally, they indicate that this is the main distinguishing feature between the two classes of PFFs as defined by the presence of O-GlcNAc and the altered structure, although the difference is stability seen in the KO neurons may also contribute.

### Altered fibril interactions could contribute to the diminished seeding activity of O-GlcNAc-modified α-Syn fibrils

We next set out to test potential molecular mechanisms to reconcile the *in vitro* and *in vivo* seeding capacity of α-Syn(gS87) PFFs. Given that the C-terminal domain of α-Syn decorates the surface of PFFs, we wondered whether cleavage of the C-terminus of internalized α-Syn(gS87) PFFs might interfere with their ability to seed the aggregation of endogenous α-Syn. To directly test this possibility *in vitro*, we investigated the effect of post-fibrillization C-terminal cleavage of PFFs on their seeding activity *in vitro*. Towards this, we generated PFFs and subjected them to different amounts of calpain and analyzed the production of the 12 kDa fragment by SDS-PAGE (Figure S10a). Using this experiment as a guide, we normalized the amounts of 12 kDa α-Syn and α-Syn(gS87) PFFs (Figure S10b) and used them to seed unmodified, human protein aggregation. After analysis by sedimentation, we observed very little difference between the full-length and 12 kDa PFFs for either unmodified or α-Syn(gS87) PFFs (Figure 6a), and we made the same observation with mouse monomeric α-Syn (Figure 6b). These results suggest that the cleavage of PFFs in neurons cannot explain the lack of seeded aggregation for α-Syn(gS87); however, they show directly for the first time to our knowledge that post-fibrilization cleave of PFFs (even unmodified ones) does not significantly influence their seeding activity *in vitro*.

**Figure 6.**
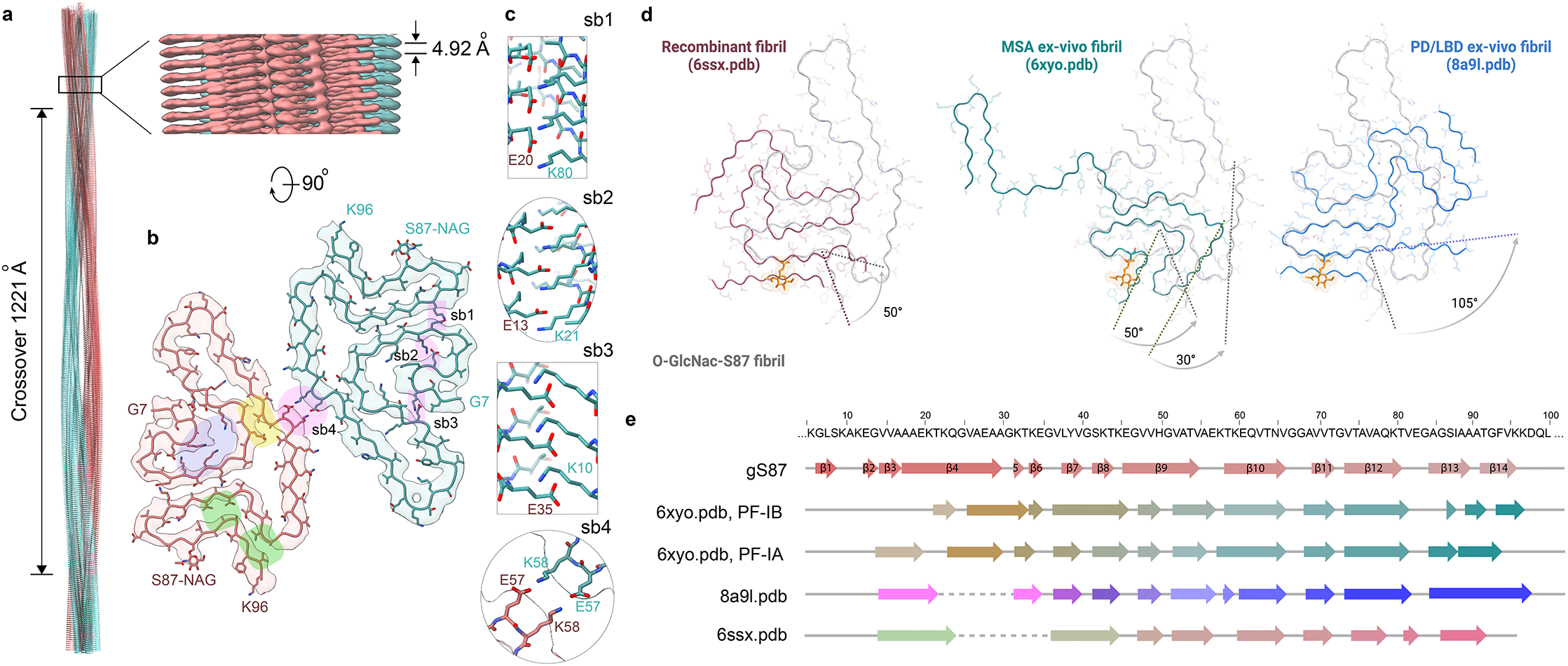
Cryo-EM structure and atomic model of the gS87 double filaments and comparison to other fibril structures. a) The cryo-EM structure of full-length gS87 fibril with a crossover distance of 1221 Å, and a close-up side view of the reconstructed 3D map showing the distance between layers, helical rise, of 4.92 Å. b) A cross-section of the density map (salmon and green) overlaid with the atomic model. Areas highlighted in pink indicate potential salt bridges. Areas highlighted in yellow and green indicate hydrophobic interdigitated interfaces. The area highlighted in purple indicates a lysine-rich region. c) Five possible salt bridges connecting the two protofilaments (sb1), and within one protofilament (sb2, sb3 and sb4). d) Superimposition of the gS87 protofilament (light grey ribbon) to an unmodified *in vitro* fibril (red ribbon, pdf: 6ssx), a MSA *ex vivo* fibril (green ribbon, pdb: 6xyo), and a PD/LBD *ex vivo* fibril (blue ribbon, pdb: 8a9l). e) Secondary structural comparison of gS87 double filaments to MSA *ex-vivo* fibril (protofilament IA, protofilament IB, pdb:6xyo) and PB/LBD *ex-vivo* fibril (pdb: 8a9l) with primary sequence indicated on the top.

Given this result, we reasoned that the lack of seeding in neurons and *in vivo* may result from altered protein interactions between the endogenous α-Syn monomers and α-Syn(gS87) PFFs. The differential interactions between unmodified and O-GlcNac modified α-Syn PFFs and chaperones could also explain their differential seeding activity. The small heat shock proteins (sHSPs)^46^ represent one such interaction that can directly inhibit PFF seeding^47^. We hypothesized that sHSPs may more potently inhibit α-Syn(gS87) PFF seeding compared to unmodified PFFs and directly tested this possibility with HSP27, an sHSP highly-expressed in the brain. Specifically, we performed seeded aggregation experiments identical to those in Figure 2, with either α-Syn or α-Syn(gS87) PFFs and unmodified monomer, in the presence of different ratios of HSP27 (Figure 6c). As expected based on our previous results^48^, HSP27 displayed partial inhibition of aggregation seeded by α-Syn PFFs at a ratio of 400:1 and full inhibition at 100:1 (α-Syn monomer:HSP27). Strikingly, HSP27 displayed more potent inhibition of seeding by α-Syn(gS87) PFFs at 400:1. These results confirm our hypothesis, at least *in vitro*, and strongly suggest that α-Syn(gS87) PFFs are more effectively chaperoned in neurons and most-likely have other different protein interactions that prevent seeded aggregation and the development of pathology *in vivo*.

### The structure of α-Syn(gS87) amyloids shows marked differences from previous *in vitro* and *ex vivo* fibrils

Finally, we set out to determine whether O-GlcNAc alters the structure of α-Syn. Towards this goal, we sought to determine the structure of α-Syn(gS87) fibrils using cryo-EM. As described above, we subjected α-Syn(gS87) to aggregation conditions, and we confirmed the presence of fibrils using TEM of negative stained grids and cryo-EM of vitrified grids (Figure S12). Both techniques revealed various fibril morphologies, including double, triple, and quadruple filaments comprised of two, three, and four protofilaments, respectively (Figure S12b). The most abundant species was the double filament, which had a diameter of approximately 20 nm. The triple and quadruple filaments had a diameter of approximately 30 nm and 40 nm, respectively. We collected cryo-EM images and manually picked all filaments for data processing and structure determination. Consistent with our previous observation, 2D classification showed three predominant fibril populations, with double filaments comprising the majority (68%). Due to the limited number of segments present in the triple and quadruple filament morphologies, we only performed 3D reconstruction with the double filaments. The 3D classification resulted in one main class with two protofilaments seemingly related by a 2-fold axis. The resulting density map, with a resolution of 4.0 Å, corresponded to a crossover of 1221 Å, an optimized twist angle of -0.7° (assumed to be left-handed), and a helical rise of 4.92 Å (Figure 6a).

Despite the limited resolution, the density map could fit a reasonable α-Syn(gS87) model comprising the residues from glycine 7 to lysine 96 and an extra density corresponding to the sugar on serine 87. A surprising observation is that the structural core of the α-Syn(gS87) fibrils is highly charged. The two protofilaments are held together by salt bridges at the interface, E57-K58 and K58-E57 (Figure 6b&c; sb1). Other salt bridges may contribute to the folding of the α-Syn(gS87) fiber-structure (Figure 6b&c; sb2, sb3, and sb4), and 5 lysine residues are packed in the center of each protofilament surrounding a small channel (Fig 6b, highlighted in blue). Like other amyloid fibrils, α-Syn(gS87) fibrils are also characterized by hydrophobic interfaces, such as the one created by residues Ala53-Ala30-Val55, Ala68-Val69-Val95, Ile88-Ala90-Val76 (Figure 2b, highlighted in yellow and green), but they are comparatively small. Possibly, as a result, α-Syn(gS87) fibrils are not likely to be particularly stable *in vitro* when compared to other amyloids, as they have a solvation energy of approximately -24.0 kcal/mol (Figure S13)^15,49^.

The structural comparison with *ex-vivo* fibrils shows that α-Syn(gS87) fibrils share some structural similarities with fibrils obtained from individuals with MSA and PD/LBD^17,50^. When compared to the structures of fibrils extracted from patients, α-Syn(gS87) fibrils share a secondary structure composition of the C-terminal end, starting in residue lysine 57 (Figure 6). However, the presence of the sugar on serine 87 makes the structure of α-Syn(gS87) fibrils incompatible with the conformation found in MSA or PD/LBD *ex-vivo* fibrils. In the case of MSA ex-vivo fibrils, the O-GlcNAc moiety would force the C-terminal conformation to rotate 30-50 ° (Figure 6d, middle). In the case of PD/LBD, the sugar moiety would be incompatible with the interface that is formed between the structure core and the additional protein density of unknown nature (Figure 6d, right). We also compared α-Syn(gS87) fibrils to the amyloid strain formed in vitro under our aggregation conditions^51^ and again found that the O-GlcNAc is incompatible with the structure (Figure 6d, left). The secondary structure composition of the compared models is detailed in Figure 6e. This structural model, although at fairly low resolution, confirms that α-Syn(gS87) fibrils are quite different from both the pathogenic unmodified fibrils we prepared here and the amyloids isolated from patient brains to date. Our results suggest that PTMs contribute to expanding the conformational landscape of amyloid fibrils and are key determinants of amyloid structural diversity. These observations point to deciphering the PTM signature of brain-derived fibrils and generation of site-specifically modified fibrils in vitro as essential first steps to bridge the gap between the structure of α-Syn fibrils produced *in vitro* and those isolated from diseased brain and to recapitulate the biochemical and structural properties of pathological aggregates in the brain.

## DISCUSSION

Our results demonstrate that O-GlcNAc can cause the formation of an alternative strain of α-Syn(gS87) amyloids with diminished *in vivo* seeding activity. Interestingly, although α-Syn(gS87) PFFs can readily seed additional aggregation *in vitro* and template their structure onto unmodified α-Syn, they display very low levels of seeding activity in neurons or *in vivo*. These observations suggest that the effect of this modification not only on the structure of the fibrils but also on how they interact with other molecules, proteins and organelles in the cellular environment is a key determinant of their seeding activity *in vivo*. We reasoned that O-GlcNAc may alter the α-Syn-fibril interactome in ways that favor interaction with chaperones or other proteins that could modify the surfaces of the fibrils and inhibit their seeding activity. To test this hypothesis, we focused on the interactions of sHSPs as an attractive inhibitory interaction. Gratifyingly, we demonstrated that HSP27, which is highly expressed in the brain, has a greater ability to inhibit seeding by α-Syn(gS87) PFFs compared to unmodified PFFs *in vitro*. We think that it is very likely that other proteins may also be responsible and plan to explore how the overall interactome might be altered using proteomics approaches in the future. Finally, we took advantage of cryo-EM to generate a model of the α-Syn(gS87) PFFs that appears to be a novel amyloid strain that is very different from the unmodified PFFs used here as well as aggregates analyzed from synucleinopathy-patient samples. Notably, the α-Syn(gS87) strain has a very small interface between the two protofilaments, an increase in charged interactions that form the core of the fiber, and hydrophobic patches on the outside of the core structure (Figure S12). Additionally, the β-sheet character of the protein is extended further into the N-terminus.

We speculate that any of these features may allow HSP27, and likely other protein factors, to recognize the amyloid core directly. Additionally, the structure is likely to change the display of the N- and C-terminal extensions or “fuzzy coat” of the protein in ways that could also be differentiated from the unmodified PFFs. Our findings also show that not all the fibrils in the brain are pathogenic and suggest that correlating fibril/ pathology load in the brain and neurodegeneration of clinical symptoms may not be the best approach to assessing the efficacy of therapeutics in clinical trials. This underscores the critical importance of developing diagnostics and therapeutics that account for not only the structural diversity of fibrils but also their differential pathogenic properties.

Taken together, we believe that these results have important implications for amyloidogenesis in general and the exploitation of O-GlcNAc in the treatment of PD and other neurodegenerative diseases. To our knowledge, this is the first example of a PTM that occurs on α-Syn monomers that can cause the formation of an amyloid strain that shows such dramatically reduced pathogenicity. Additionally, the divergence between our *in vitro* and cellular seeding results is striking. These data suggest that interactions between α-Syn fibrils and other cellular proteins or organelles are key determinants of their pathology formation and toxicity. Furthermore, they show that connections between the *in vitro* seeding and pathogenic potential of different amyloids might be decoupled and should be considered in future experiments. Our results also support the continued development of OGA inhibitors to treat PD, as increased O-GlcNAc has the strong potential to both slow the initial aggregation of α-Syn monomers but also result in the formation of *in vivo* seeding-incompetent fibrils. This is particularly true given the inconsistent results of recent antibody therapies that target the amyloid aggregates. Two compelling hypotheses for the failure of some of these drugs are that loss of the monomeric protein and/or the aggregation process overall are the underlying toxic events. The results here, combined with our previous publications, provide a strong foundation that increased O-GlcNAc will maintain the levels of soluble α-Syn and slow the aggregation process through the direct inhibition of the initiation of aggregation^34,35^ and reduced seeding (Figure 5c). Finally, we think that it would be interesting to examine the consequences of O-GlcNAc on Tau amyloid structures more closely, as the major site of O-GlcNAc (S400) also slows but does not completely block its aggregation^33^.

## Supporting information

Experimental Methods

## ACKNOWLEDGMENTS

M.R.P acknowledges support from the Michael J. Fox Foundation for Parkinson’s Research (MJFF-008121), the Anton Burg Foundation, and the National Institutes of Health (R01GM114537). A-L.M-M. and H.A.L. acknowledge support from the École Polytechnique Fédérale de Lausanne. V.M.-Y.L. and K.C.L. acknowledge support from the National Institutes of Health (U19AG062418). L.S. acknowledges support from the Department of Energy, Laboratory Directed Research and Development program at SLAC National Accelerator Laboratory (DE-AC02-76SF00515). A.T.B. was supported as a Dornsife Chemistry-Biology Interface Trainee. The Electron Microscopy Facility at UTSW is supported by the Cancer Prevention & Research Institute of Texas (RP170644) and the National Institutes of Health (1S10OD021685-01A1, 1S10OD020103-01). TEM images were collected at the USC Core Center of Excellence in Nano Imaging, and qTOF analysis was performed at the USC Chemistry Mass Spectrometry Core. Some computational resources used for structure determination were provided by the BioHPC supercomputing facility located in the Lyda Hill Department of Bioinformatics at UTSW.

## AUTHOR CONTRIBUTIONS

A.T.B., A-L.M-M., B.A.N., V.M.-Y.L., K.C.L, L.S., H.A.L. and M.R.P. designed experiments and interpreted data. A.T.B. carried out protein synthesis and characterization, *in vitro* aggregation and seeding experiments, and analysis of fibril phosphorylation and O-GlcNAc modification. A-L.M-M. performed cellular analysis of PFFs: uptake, processing, stability, pathogenicity, aggregation, and co-localization. B.A.N. performed structure experiments: fibril formation, screening, data collection and processing, and modeling. M.H. carried out cellular and *in vivo* pathogenicity experiments: protein aggregation, neuron treatment, mouse injections, staining, and data collection and analysis. A.J. and E.R.D performed HSP27 inhibition experiments. Y.J. carried out protein aggregation for cellular analysis and characterization of fibrils before and after sonication. P.S. and V.S. performed particle picking for cryo-EM analysis. S.A. and R.P. carried out screening for cryo-EM experiments. L.S. carried out structural modeling. A.T.B., A-L.M-M., B.A.N., M.H., Y.J., K.C.L, L.S., H.A.L. and M.R.P. prepared figures. A.T.B., A-L.M-M., B.A.N., L.S., H.A.L. and M.R.P. wrote the manuscript.

## REFERENCES

1. Eisenberg, D. & Jucker, M. The Amyloid State of Proteins in Human Diseases. Cell 148, 1188–1203 (2012).

2. Dobson, C. M., Knowles, T. P. J. & Vendruscolo, M. The Amyloid Phenomenon and Its Significance in Biology and Medicine. Csh Perspect Biol 12, a033878 (2020).

3. Willbold, D., Strodel, B., Schröder, G. F., Hoyer, W. & Heise, H. Amyloid-type Protein Aggregation and Prion-like Properties of Amyloids. Chem Rev 121, 8285–8307 (2021).

4. Cascella, R., Bigi, A., Cremades, N. & Cecchi, C. Effects of oligomer toxicity, fibril toxicity and fibril spreading in synucleinopathies. Cell Mol Life Sci 79, 174 (2022).

5. Volpicelli-Daley, L. A. et al. Exogenous α-Synuclein Fibrils Induce Lewy Body Pathology Leading to Synaptic Dysfunction and Neuron Death. Neuron 72, 57–71 (2011).

6. Mahul-Mellier, A.-L. et al. The making of a Lewy body: the role of α-synuclein post-fibrillization modifications in regulating the formation and the maturation of pathological inclusions. Biorxiv 500058 (2018) doi:10.1101/500058.

7. Mahul-Mellier, A.-L. et al. The process of Lewy body formation, rather than simply α-synuclein fibrillization, is one of the major drivers of neurodegeneration. Proc National Acad Sci 117, 4971–4982 (2020).

8. Frost, B., Jacks, R. L. & Diamond, M. I. Propagation of Tau Misfolding from the Outside to the Inside of a Cell*. J Biol Chem 284, 12845–12852 (2009).

9. Guo, J. L. & Lee, V. M.-Y. Seeding of Normal Tau by Pathological Tau Conformers Drives Pathogenesis of Alzheimer-like Tangles*. J Biol Chem 286, 15317–15331 (2011).

10. Kaufman, S. K. et al. Tau Prion Strains Dictate Patterns of Cell Pathology, Progression Rate, and Regional Vulnerability In Vivo. Neuron 92, 796–812 (2016).

11. Luk, K. C. et al. Pathological α-synuclein transmission initiates Parkinson-like neurodegeneration in nontransgenic mice. Science (New York, NY) 338, 949–953 (2012).

12. Luk, K. C. et al. Molecular and Biological Compatibility with Host Alpha-Synuclein Influences Fibril Pathogenicity. Cell Reports 16, 3373–3387 (2016).

13. Gracia, P., Camino, J. D., Volpicelli-Daley, L. & Cremades, N. Multiplicity of α-Synuclein Aggregated Species and Their Possible Roles in Disease. Int J Mol Sci 21, 8043 (2020).

14. Shi, Y. et al. Structure-based classification of tauopathies. Nature 598, 359–363 (2021).

15. Kametani, F. & Hasegawa, M. Structures of tau and α-synuclein filaments from brains of patients with neurodegenerative diseases. Neurochem Int 158, 105362 (2022).

16. Shahnawaz, M. et al. Discriminating α-synuclein strains in Parkinson’s disease and multiple system atrophy. Nature 578, 273–277 (2020).

17. Schweighauser, M. et al. Structures of α-synuclein filaments from multiple system atrophy. Nature 585, 464–469 (2020).

18. Perren, A. V. der et al. The structural differences between patient-derived α-synuclein strains dictate characteristics of Parkinson’s disease, multiple system atrophy and dementia with Lewy bodies. Acta Neuropathol 139, 977–1000 (2020).

19. Lloyd, G. M. et al. Unique seeding profiles and prion-like propagation of synucleinopathies are highly dependent on the host in human α-synuclein transgenic mice. Acta Neuropathol 1–23 (2022) doi:10.1007/s00401-022-02425-4.

20. Oueslati, A., Fournier, M. & Lashuel, H. A. Chapter 7 Role of post-translational modifications in modulating the structure, function and toxicity of α-synuclein Implications for Parkinson’s disease pathogenesis and therapies. Prog Brain Res 183, 115–145 (2010).

21. Schmid, A. W., Fauvet, B., Moniatte, M. & Lashuel, H. A. Alpha-synuclein Post-translational Modifications as Potential Biomarkers for Parkinson Disease and Other Synucleinopathies. Mol Cell Proteomics 12, 3543– 3558 (2013).

22. Wesseling, H. et al. Tau PTM Profiles Identify Patient Heterogeneity and Stages of Alzheimer’s Disease. Cell 183, 1699-1713.e13 (2020).

23. Limorenko, G. & Lashuel, H. A. To target Tau pathologies, we must embrace and reconstruct their complexities. Neurobiol Dis 161, 105536 (2021).

24. Pancoe, S. X. et al. Effects of Mutations and Post-Translational Modifications on α-Synuclein In Vitro Aggregation. J Mol Biol 434, 167859 (2022).

25. Ma, J., Wu, C. & Hart, G. W. Analytical and Biochemical Perspectives of Protein O-GlcNAcylation. Chem Rev 121, 1513–1581 (2021).

26. Lee, B. E., Suh, P.-G. & Kim, J.-I. O-GlcNAcylation in health and neurodegenerative diseases. Exp Mol Medicine 53, 1674–1682 (2021).

27. Balana, A. T. & Pratt, M. R. Mechanistic roles for altered O-GlcNAcylation in neurodegenerative disorders. Biochem J 478, 2733–2758 (2021).

28. Liu, F. et al. Reduced O-GlcNAcylation links lower brain glucose metabolism and tau pathology in Alzheimer’s disease. Brain 132, 1820–1832 (2009).

29. Wang, A. C., Jensen, E. H., Rexach, J. E., Vinters, H. V. & Hsieh-Wilson, L. C. Loss of O-GlcNAc glycosylation in forebrain excitatory neurons induces neurodegeneration. Proceedings of the National Academy of Sciences of the United States of America 113, 15120–15125 (2016).

30. Bartolomé-Nebreda, J. M., Trabanco, A. A., Velter, A. I. & Buijnsters, P. O-GlcNAcase inhibitors as potential therapeutics for the treatment of Alzheimer’s disease and related tauopathies: analysis of the patent literature. Expert Opin Ther Pat 31, 1117–1154 (2021).

31. Yuzwa, S. A. et al. Increasing O-GlcNAc slows neurodegeneration and stabilizes tau against aggregation. Nat Chem Biol 8, 393–399 (2012).

32. Permanne, B. et al. O-GlcNAcase Inhibitor ASN90 is a Multimodal Drug Candidate for Tau and α-Synuclein Proteinopathies. Acs Chem Neurosci 13, 1296–1314 (2022).

33. Yuzwa, S. A., Cheung, A. H., Okon, M., McIntosh, L. P. & Vocadlo, D. J. O-GlcNAc Modification of tau Directly Inhibits Its Aggregation without Perturbing the Conformational Properties of tau Monomers. J Mol Biol 426, 1736–1752 (2014).

34. Marotta, N. P. et al. O-GlcNAc modification blocks the aggregation and toxicity of the protein α-synuclein associated with Parkinson’s disease. Nat Chem 7, 913–920 (2015).

35. Levine, P. M. et al. α-Synuclein O-GlcNAcylation alters aggregation and toxicity, revealing certain residues as potential inhibitors of Parkinson’s disease. Proc National Acad Sci 116, 201808845 (2019).

36. Thompson, R. E. & Muir, T. W. Chemoenzymatic Semisynthesis of Proteins. Chem Rev 120, 3051–3126 (2019).

37. Moon, S. P., Javed, A., Hard, E. R. & Pratt, M. R. Methods for Studying Site-Specific O-GlcNAc Modifications: Successes, Limitations, and Important Future Goals. Jacs Au 2, 74–83 (2021).

38. Polinski, N. K. et al. Best Practices for Generating and Using Alpha-Synuclein Pre-Formed Fibrils to Model Parkinson’s Disease in Rodents. J Park Dis Preprint, 1–20 (2018).

39. Kumar, S. T., Donzelli, S., Chiki, A., Syed, M. M. K. & Lashuel, H. A. A simple, versatile and robust centrifugation‐based filtration protocol for the isolation and quantification of α‐synuclein monomers, oligomers and fibrils: Towards improving experimental reproducibility in α‐synuclein research. J Neurochem 153, 103–119 (2020).

40. Bousset, L. et al. Structural and functional characterization of two alpha-synuclein strains. Nat Commun 4, 2575 (2013).

41. Giorgi, F. D. et al. Novel self-replicating α-synuclein polymorphs that escape ThT monitoring can spontaneously emerge and acutely spread in neurons. Sci Adv 6, eabc4364 (2020).

42. Kumar, S. T. et al. A NAC domain mutation (E83Q) unlocks the pathogenicity of human alpha-synuclein and recapitulates its pathological diversity. Sci Adv 8, eabn0044 (2022).

43. Fares, M. B., Jagannath, S. & Lashuel, H. A. Reverse engineering Lewy bodies: how far have we come and how far can we go? Nat Rev Neurosci 22, 111–131 (2021).

44. Mishizen‐Eberz, A. J. et al. Distinct cleavage patterns of normal and pathologic forms of α‐synuclein by calpain I in vitro. J Neurochem 86, 836–847 (2003).

45. Karpowicz, R. J. et al. Selective imaging of internalized proteopathic α-synuclein seeds in primary neurons reveals mechanistic insight into transmission of synucleinopathies. J Biol Chem 292, 13482–13497 (2017).

46. Hartl, F. U., Bracher, A. & Hayer-Hartl, M. Molecular chaperones in protein folding and proteostasis. Nature 475, 324–332 (2011).

47. Cox, D. et al. The small heat shock protein Hsp27 binds α-synuclein fibrils, preventing elongation and cytotoxicity. J Biol Chem 293, 4486–4497 (2018).

48. Balana, A. T. et al. O-GlcNAc modification of small heat shock proteins enhances their anti-amyloid chaperone activity. Nat Chem 1–10 (2021) doi:10.1038/s41557-021-00648-8.

49. Nguyen, B. A. et al. Structural polymorphism of amyloid fibrils in cardiac transthyretin amyloidosis revealed by cryo-electron microscopy. Biorxiv 2022.06.21.496949 (2022) doi:10.1101/2022.06.21.496949.

50. Yang, Y. et al. Cryo-EM structures of α-synuclein filaments from Parkinson’s disease and dementia with Lewy bodies. Biorxiv 2022.07.12.499706 (2022) doi:10.1101/2022.07.12.499706.

51. Guerrero-Ferreira, R. et al. Two new polymorphic structures of human full-length alpha-synuclein fibrils solved by cryo-electron microscopy. Elife 8, e48907 (2019).

