## Supplementary Data for "O-GlcNAc modification forces the formation of an α-Synuclein amyloid-strain with notably diminished seeding activity and pathology"

### Table of contents:

**Supplementary Figure 1.** Full size blots of seeded aggregation of human α-Syn. **Page S2**
**Supplementary Figure 2.** Full size blots of seeded aggregation of mouse α-Syn. **Page S3**
**Supplementary Figure 3.** The O-GlcNAc on α-Syn(gS87) fibers is stable to enzymatic removal. **Page S4**
**Supplementary Figure 4.** α-Syn(gS87) PFFs have diminished pathology in the amygdala and motor cortex without loss of dopaminergic neurons. **Page S5**
**Supplementary Figure 5.** Characterization of PFFs. **Page S6**
**Supplementary Figure 6.** Unmodified and α-Syn(gS87) PFFs display similar processing in neurons. **Page S7**
**Supplementary Figure 7.** Unmodified and α-Syn(gS87) PFFs display similar stability in neurons. **Page S8**
**Supplementary Figure 8.** α-Syn(gS87) PFFs have dramatically reduced seeding capacity in neurons. **Page S8**
**Supplementary Figure 9.** Aggregates that form from α-Syn(gS87) PFFs are notably reduced but display Lewy body hallmarks. **Page S9**
**Supplementary Figure 10.** Generation of truncated α-Syn PFFs. **Page S10**
**Supplementary Figure 11.** Altered protein interactions rather than inherent seeding-capacity likely explain the divergence between α-Syn(gS87) PFFs *in vitro* and *in vivo*. **Page S11**
**Supplementary Figure 12.** Cryo-EM data collection and processing of α-Syn(gS87) fibrils. **Page S12**
**Supplementary Figure 13.** Residues composition (left) and solvation energy map (right) of the α-Syn(gS87) double filaments. **Page S12**
**Supplementary Table S1.** Cryo-EM data collection, reconstruction, model building, and validation. **Page S13**

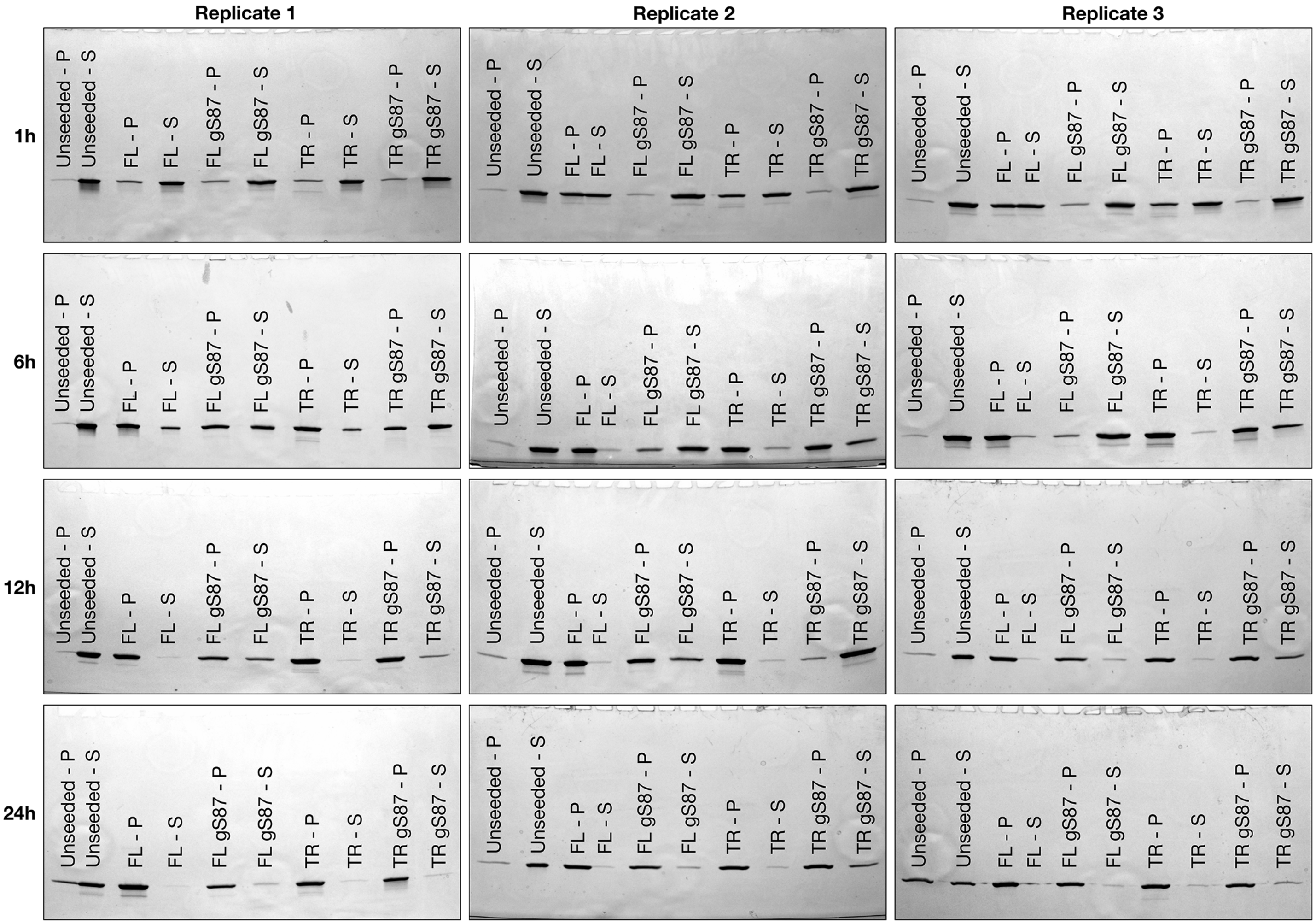


**Supplementary Figure 1. Full size blots of seeded aggregation of human α-Syn.** Raw data for Figures 2f and 6a. FL = full length α-Syn, FL gS87 = full length α-Syn(gS87), TR = truncated α-Syn, TR gS87 = truncated α-Syn(gS87), P = pellet, S = soluble.

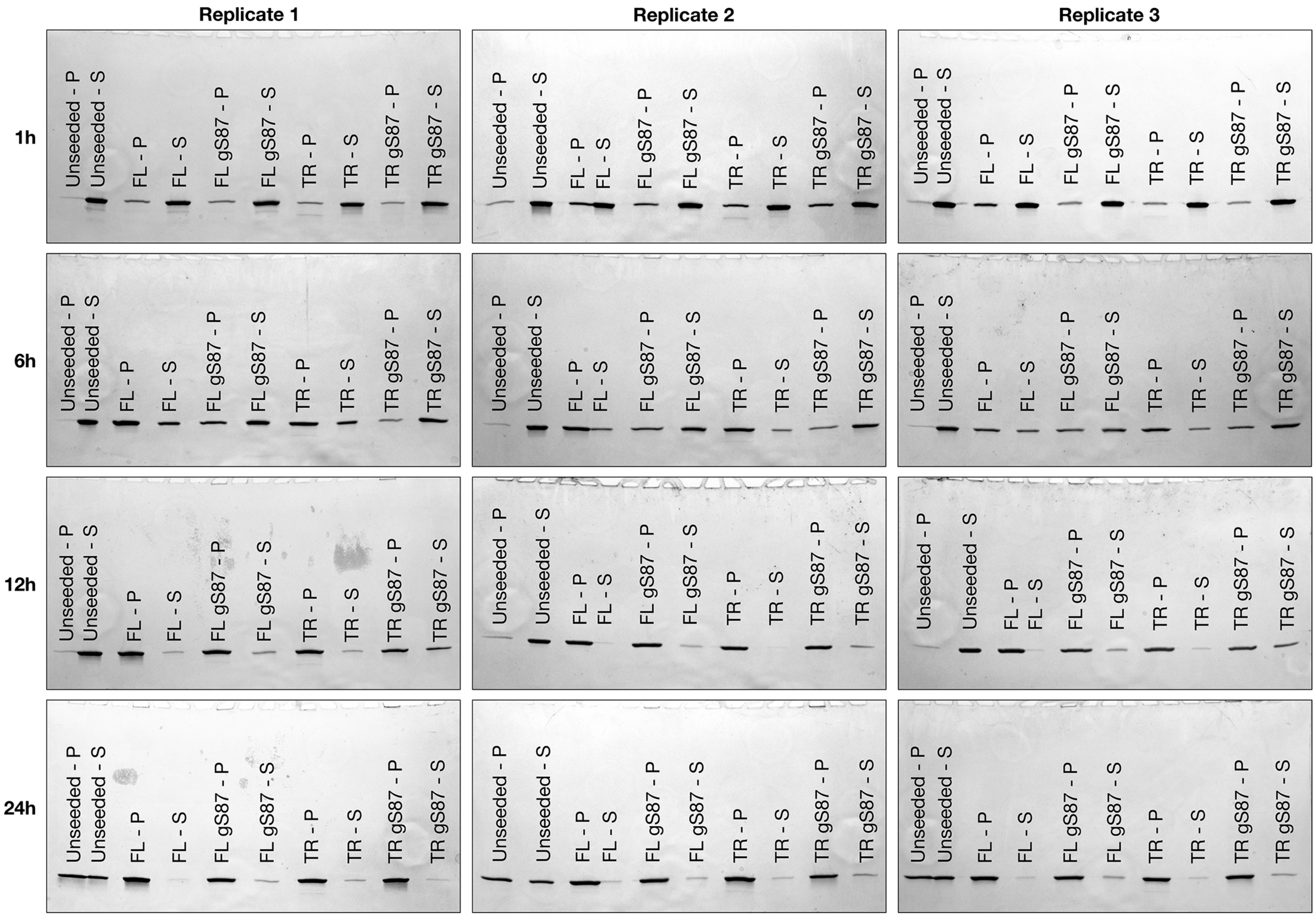


**Supplementary Figure 2. Full size blots of seeded aggregation of mouse α-Syn.** Raw data for Figures 2h and 6b. FL = full length α-Syn, FL gS87 = full length α-Syn(gS87), TR = truncated α-Syn, TR gS87 = truncated α-Syn(gS87), P = pellet, S = soluble.

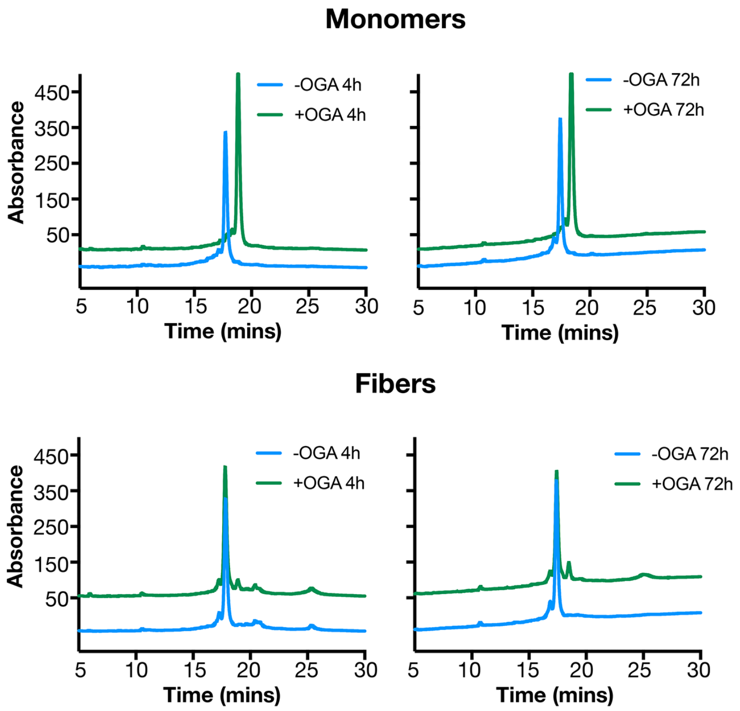


**Supplementary Figure 3. The O-GlcNAc on α-Syn(gS87) fibers is stable to enzymatic removal.** α-Syn(gS87) monomers or PFFs (25 uM) were incubated with bacterial O-GlcNAc hydrolase BtGH84 (1 uM) for the indicated lengths of time at 37°C in phosphate buffered saline. The reactions were solubilized in 8M urea and analyzed by RP-HPLC. The identity of the O-GlcNAc modified and deglycosylated proteins were confirmed by ESI-MS.

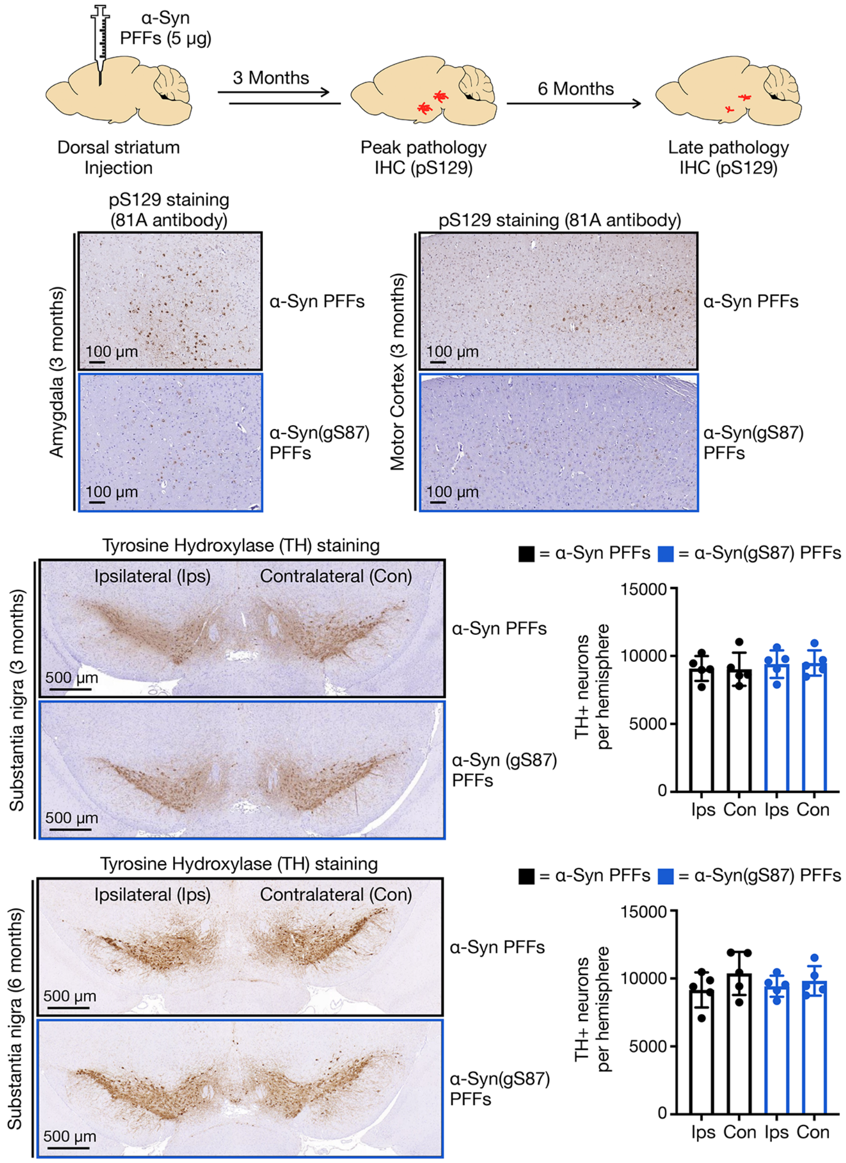


**Supplementary Figure 4. α-Syn(gS87) PFFs have diminished pathology in the amygdala and motor cortex without loss of dopaminergic neurons.** Wild-type mice were injected with α-Syn or α-Syn(gS87) PFFs (5 µg) in a single unilateral injection into the dorsal striatum. Pathology was visualized by immunohistochemistry against pS129 at 3 months post-injection. Loss of dopaminergic neurons was visualized by immunohistochemistry against tyrosine hyroxlyase (TH). Results are mean ±SEM of biological replicates (n=5). Statistical significance was determined using a Paired Student's T-test, and no statistical differences between ipsilateral and contralateral sides were found.

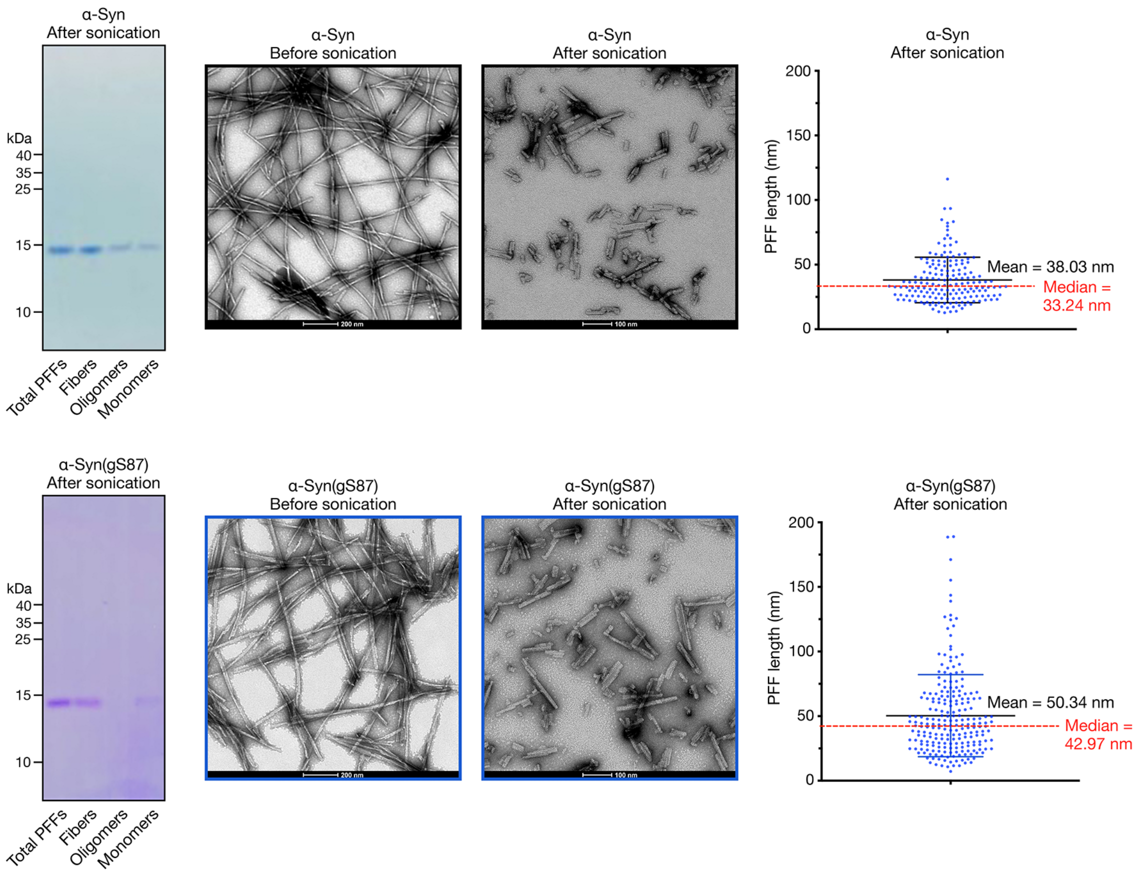


**Supplementary Figure 5. Characterization of PFFs.** α-Syn or α-Syn(gS87) were subjected to aggregation conditions (172 μM, 7d). Fibers, oligomers, and monomers contained in these aggregation reactions were then separated [Kumar et al. *J Neurochem* 153, 103–119 (2020)]. The overall aggregation reaction (Total PFFs) and the individual components were analyzed by Coomassie staining. The corresponding PFFs were also visualized before and after sonication and measured after sonication using transmission electron microscopy (TEM).

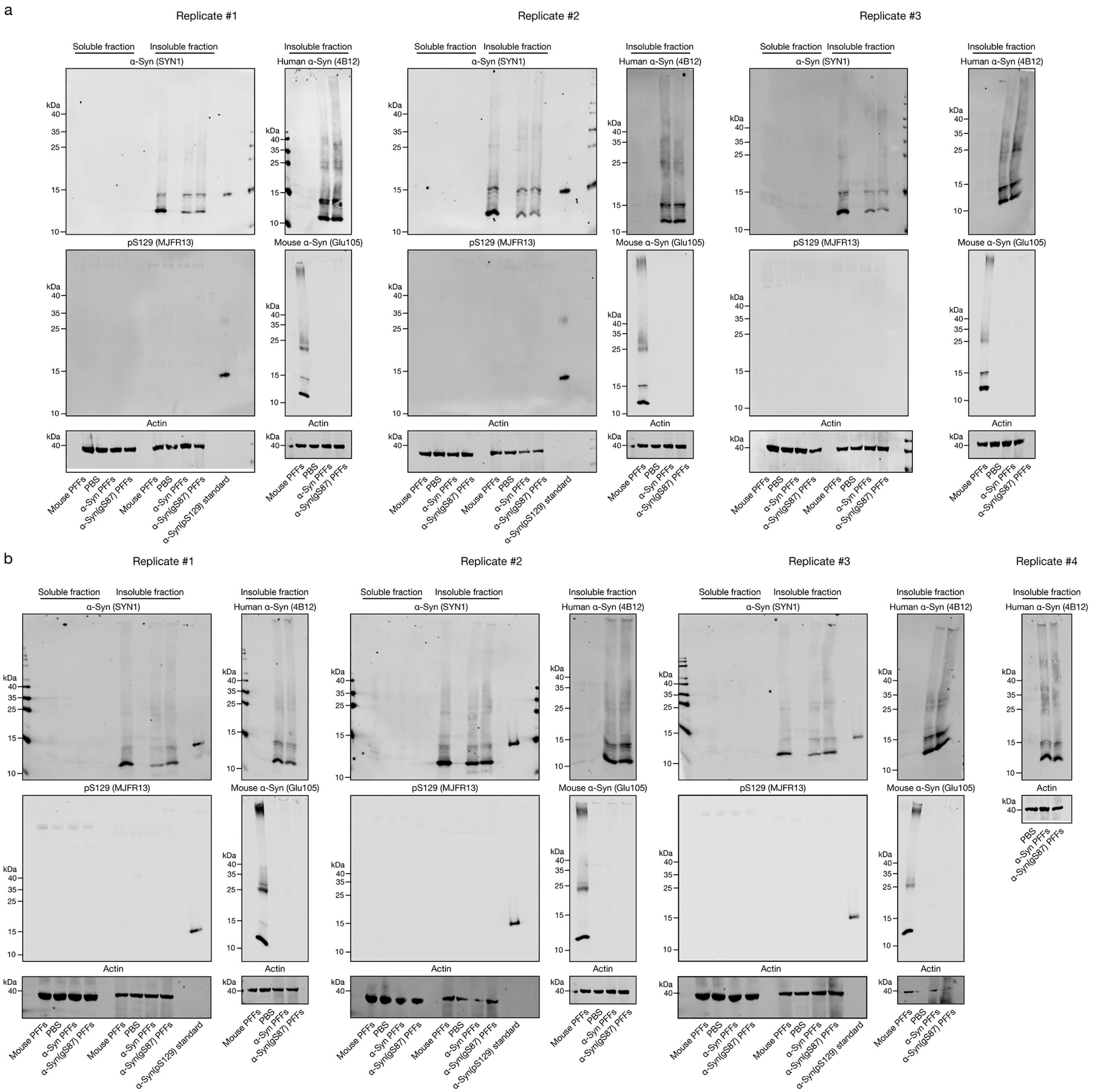


**Supplementary Figure 6. Unmodified and α-Syn(gS87) PFFs display similar processing in neurons.** Primary hippocamal neurons from α-synuclein knockout-mice at 7 days in vitro (7 DIV) were treated with the indicated human or mouse PFFs (70 nM) or PBS for different lengths of time before the following analyses. a) O-GlcNAc at S87 largely does not affect the the internalization, C-terminal cleavage to ~12 kDa fragment, or phosphorylation at S129 (pS129) of PFFs as visualized by western blotting after 14 h of treatment. b) O-GlcNAc at S87 largely does not affect the the internalization, C-terminal cleavage to ~12 kDa fragment, or phosphorylation at S129 (pS129) of PFFs as visualized by western blotting after 24 h of treatment.

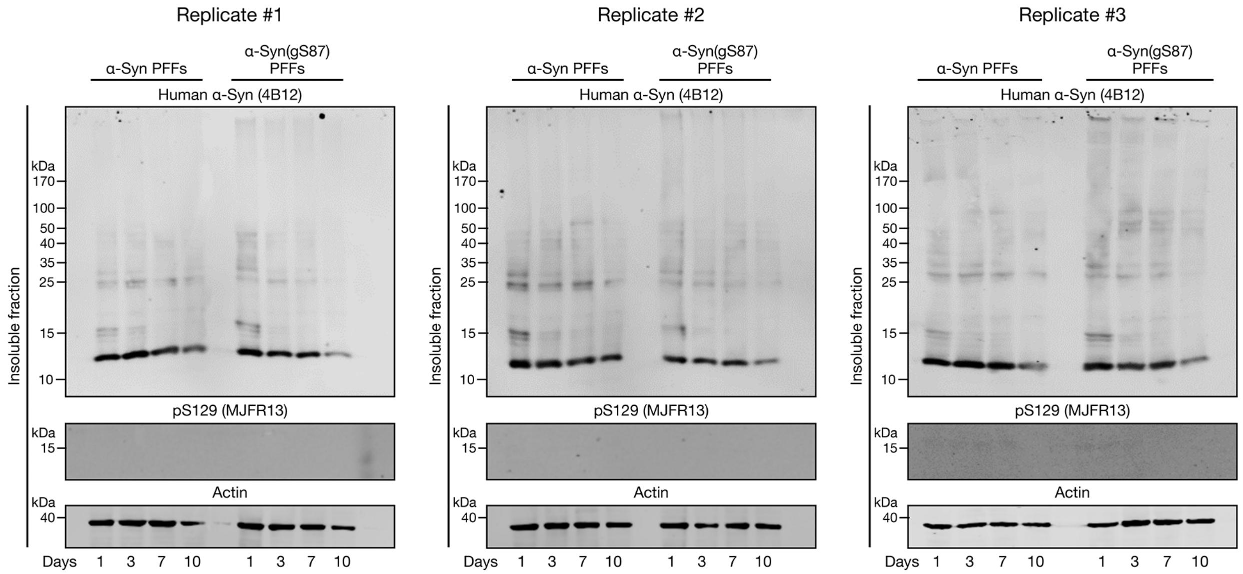


**Supplementary Figure 7. Unmodified and α-Syn(gS87) PFFs display similar stability in neurons.** Primary hippocamal neurons from α-synuclein knockout-mice at 7 days in vitro (7 DIV) were treated with the indicated human PFFs (70 nM) or PBS for different lengths of time before visualization by western blotting over 10 days of treatment.

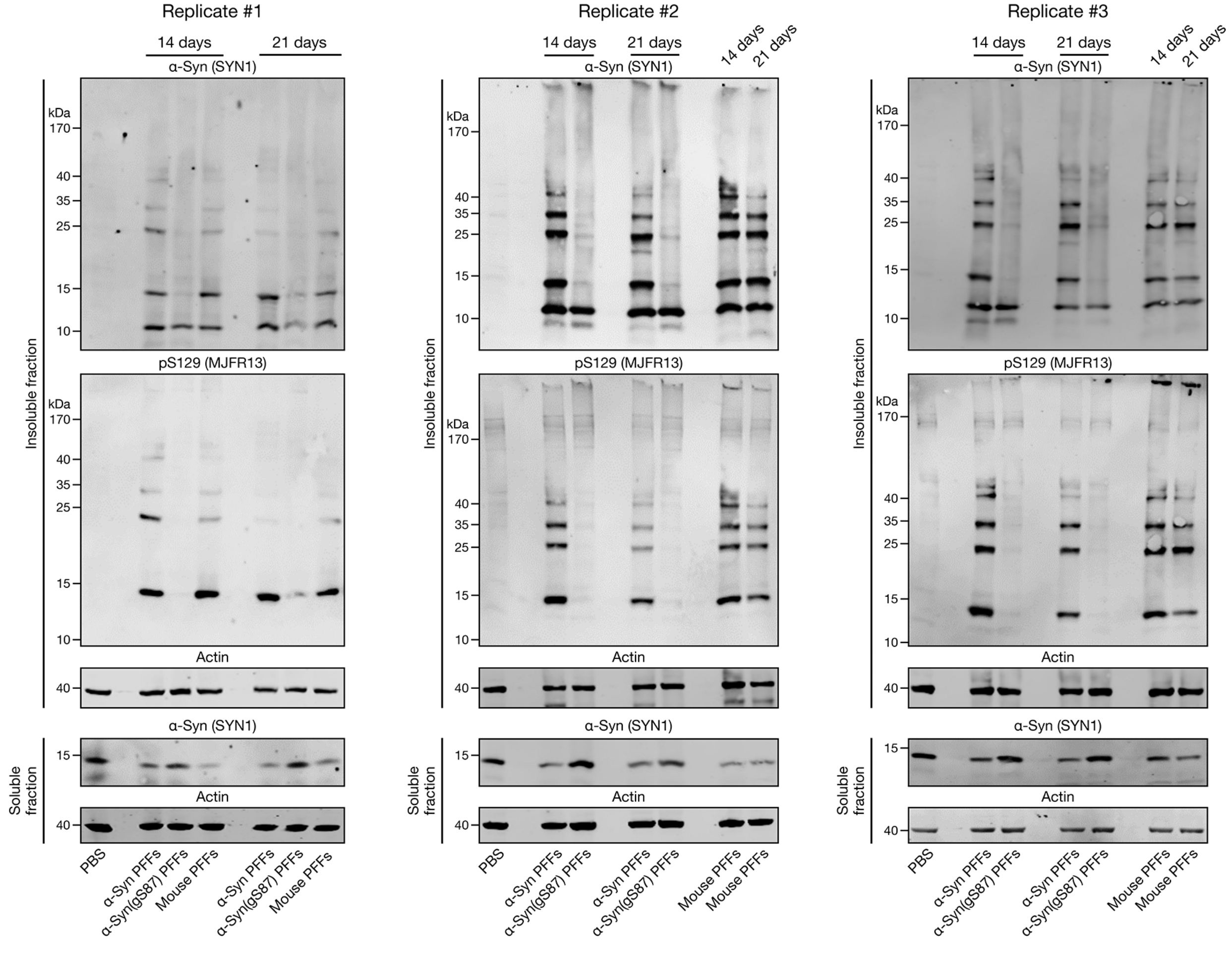


**Supplementary Figure 8. α-Syn(gS87) PFFs have dramatically reduced seeding capacity in neurons.** Primary hippocampal neurons from wild-type mice at 7 days in vitro (7 DIV) were treated with the indicated PFFs (70 nM) or PBS for different lengths of time before analysis by western blotting. Unmodified PFFs seed the aggregation of endogenous α-synuclein into insoluble and pS129-positive higher molecular-weight aggregates. O-GlcNAc at S87 dramatically reduced this seeded aggregation and more endogenous α-synuclein remained soluble.

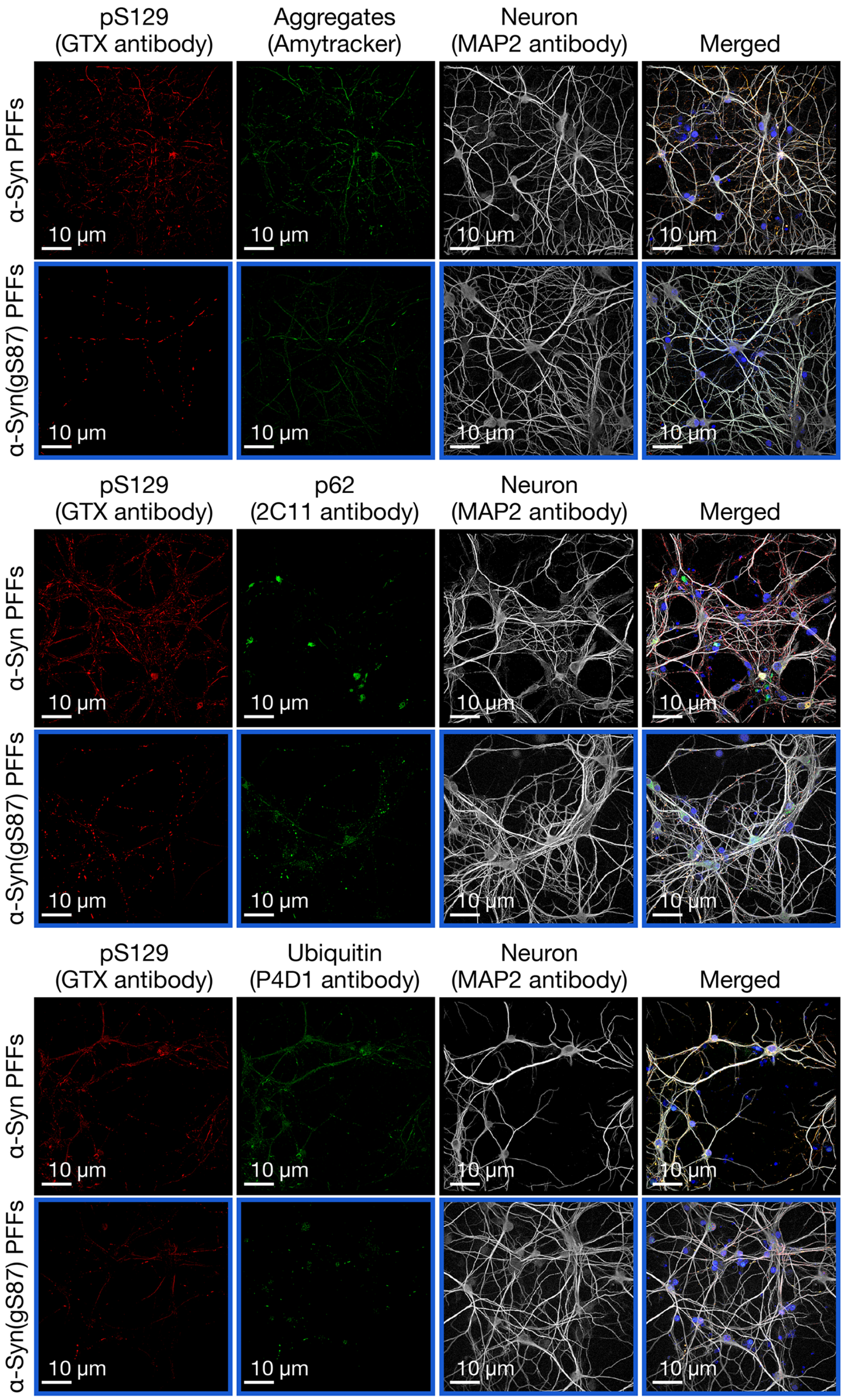


**Supplementary Figure 9. Aggregates that form from α-Syn(gS87) PFFs are notably reduced but display Lewy body hallmarks.** Primary hippocampal neurons from wild-type mice at 7 days in vitro (7 DIV) were treated with the indicated PFFs (70 nM) or PBS for 14 d before visualization of aggregate markers (amyloid, p62 & ubiquitination) by immunocytochemistry (ICC).

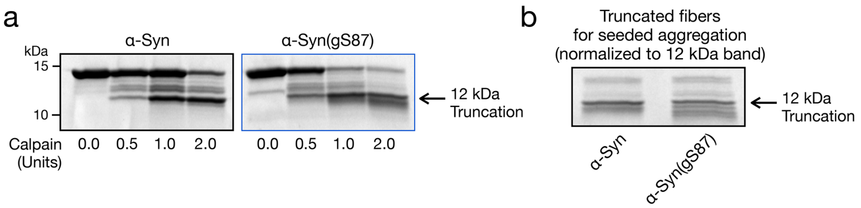


**Supplementary Figure 10. Generation of truncated α-Syn PFFs.** a) α-Syn or α-Syn(gS87) were subjected to aggregation conditions (172 μM, 7 d) before incubation with the indicated amounts of calpain and analysis by SDS-PAGE and Coomassie staining. b) The levels of PFFs truncated by 2.0 units of calpain were normalized by Coomassie staining for use in seeded aggregation experiments.

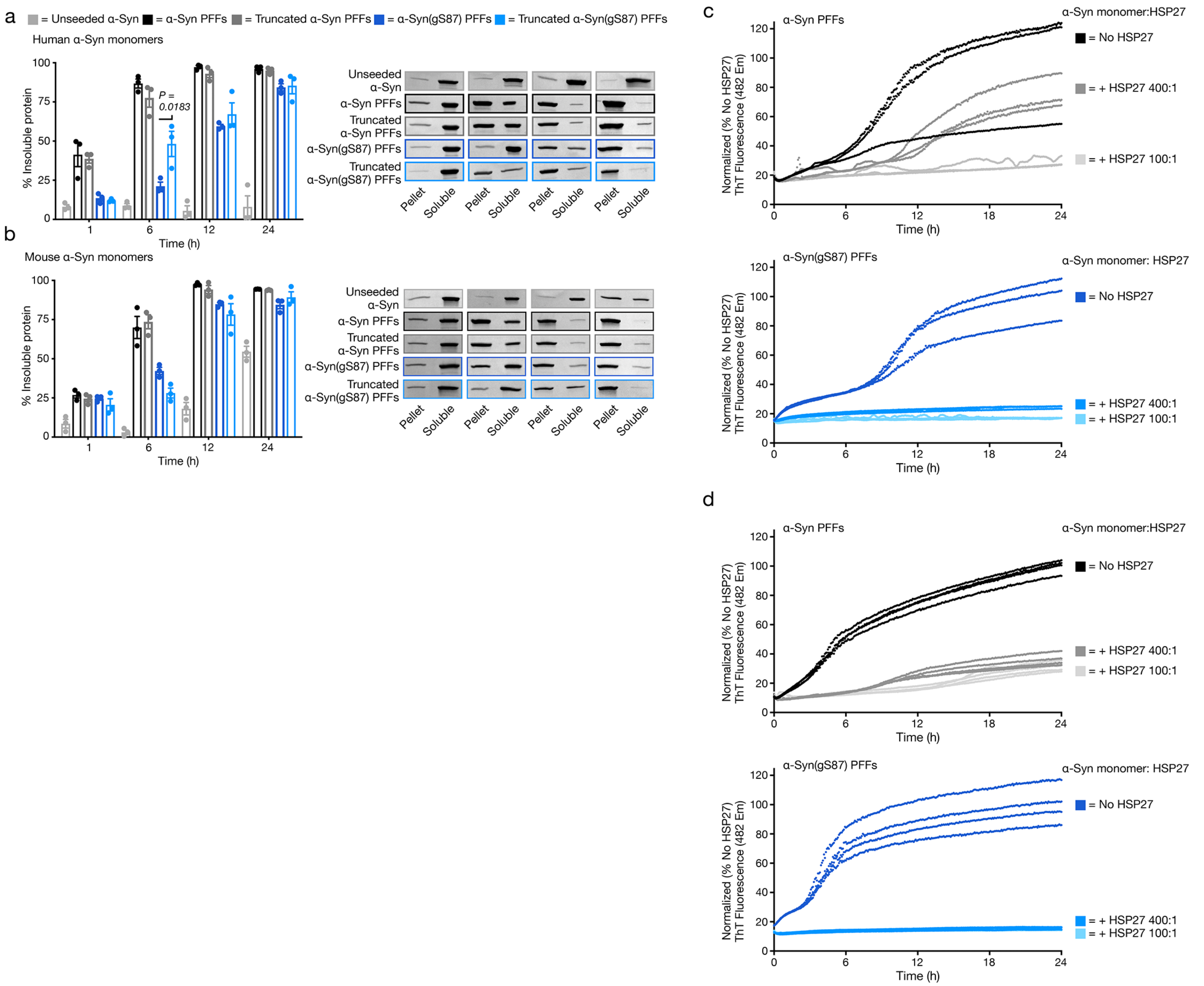


**Supplementary Figure 11. Altered protein interactions rather than inherent seeding-capacity likely explain the divergence between α-Syn(gS87) PFFs *in vitro* and *in vivo*.** a) Truncation does not dramatically alter the seeding of human α-synuclein aggregation by unmodified or α-Syn(gS87) PFFs can seed aggregation of unmodified, human α-synuclein. α-Syn PFFs (unmodified or gS87) were generated by aggregation of unmodified protein (172 μM) followed by sonication. The corresponding truncated PFFs were generated by digestion enzymatic digestion of PFFs with calpain (Figure S10). The resulting PFFs were then added to buffer or unmodified, human α-Syn (50 μM monomer concentration, 5% PFF) before aggregation and analysis by sedimentation and Coomassie staining. Results are mean ±SEM of experimental replicates (n=3). Statistical significance was determined using a one-way ANOVA test followed by Tukey's multiple comparison test. b) PFFs were generated as in (a) were then added to buffer or unmodified, mouse α-synuclein (50 μM monomer concentration, 5% PFF) before aggregation and analysis by sedimentation and Coomassie staining. Results are mean ±SEM of experimental replicates (n=3). c) HSP27 more potently inhibits aggregation seeded by α-Syn(gS87) PFFs. α-Syn monomers (50 μM) and the indicated ratios of HSP27 were mixed with the indicated PFFs (2.5 μM, 5%). The reactions were placed in a plate reader and aggregation was detected by ThT fluorescence ($\lambda_{ex}$ = 450 nm, $\lambda_{em}$ = 482 nm). d) Replicate experiment of (c).

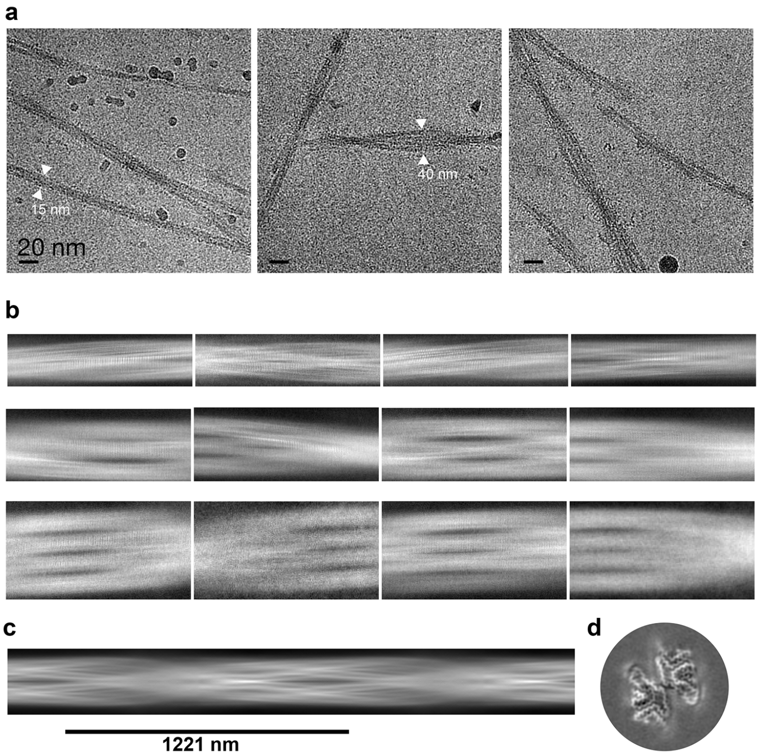


**Supplementary Figure 12. Cryo-EM data collection and processing of α-Syn(gS87) fibrils.** a) Representative cryo-EM micrographs showing polymorphisms of α-Syn(gS87) fibrils and their estimated width. Scale, 200 nm. b) Representative 2D class averages of different morphologies, double filaments – top row, triple filaments – middle row and quadruple filaments. c) Initial model of α-Syn(gS87) fibril generated by Relion3.1. d) 3D class average of α-Syn(gS87) fibril.

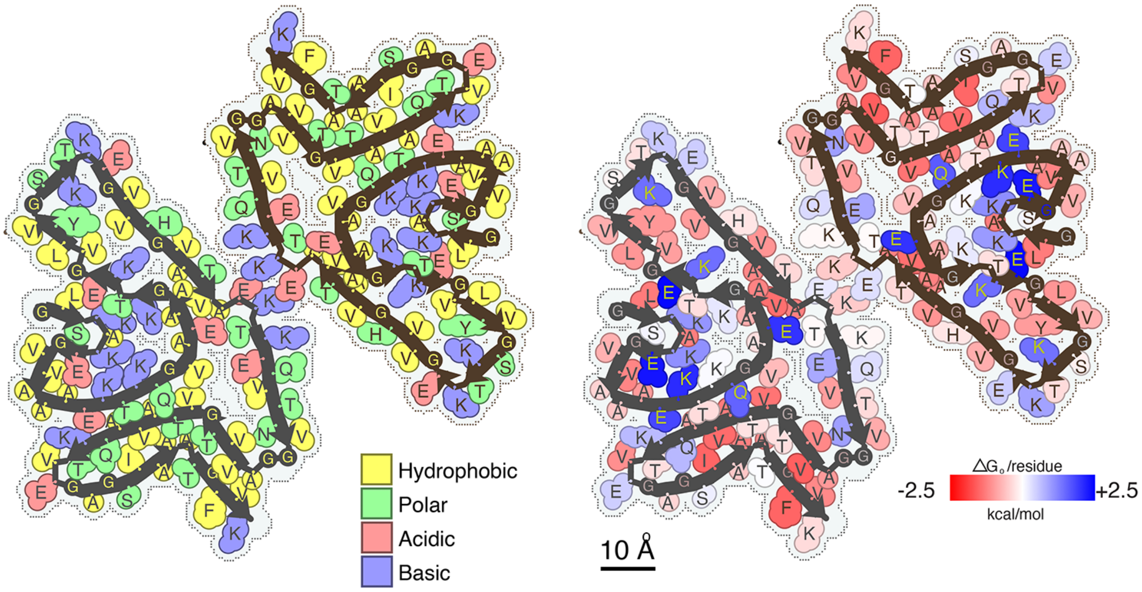


**Supplementary Figure 13. Residues composition (left) and solvation energy map (right) of the α-Syn(gS87) double filaments.** Residues on the left are colored by hydrophobicity (yellow: hydrophobic, green: polar, red: acidic, and blue: basic). Residues on the right are colored from favorable (red, -2.5 kcal/mol) to unfavorable stabilization energy (blue, 2.5 kcal.mol).

**Supplementary Table S1. Cryo-EM data collection, reconstruction, model building, and validation.**

| | |
|---|---|
| **Data collection** | |
| Microscope | Titan Krios (G3i) |
| Acceleration Voltage (kV) | 300 |
| Detector | K3 |
| Software | EPU |
| Magnification | 105,000x |
| Pixel size at detector (Å/px) | 0.86 |
| Defocus range (μm) | -0.8 to -2.2 |
| Total dose (e) | 40 |
| Exposure time (sec) | 3.6 |
| Number of movie frames | 40 |
| Usable micrograph | 7270 |
| **Helical Reconstruction** | |
| Box size (pixel) | 320 |
| Number of segments after 2D | 29633 |
| Number of segments after 3D | 22042 |
| Symmetry imposed | C1 |
| Helical rise (Å) | 4.926 |
| Helical twist (°) | -0.7 |
| Crossover length (Å) | 1221 |
| B factor | -120.12 |
| Map resolution (Å; FSC=0.143) | 4.4 |
| Map resolution (Å; FSC=0.5) | 4.7 |
| **Model composition and refinement** | |
| Non-hydrogen atoms | 3690 |
| Protein residues | 540 |
| Number of chains | 6 |
| Water/ligands | 0/0 |
| MolProbity score | 2.13 |
| Clash score | 11.45 |
| Rotamer outliers (%) | 0.00 |
| R.M.S deviations bonds (Å) | 0.002 |
| R.M.S deviations angle (°) | 0.585 |
| Ramachandran plot<br>Favored<br>Allowed<br>Outliers | <br>89.77<br>10.23<br>0 |
| CaBLAM outliers (%) | 4.65 |
| Model vs Data | 0.62 |
