## Supplementary material for "O-GlcNAc modification forces the formation of an α-Synuclein amyloid-strain with notably diminished seeding activity and pathology": Experimental Methods

**General.**

Solvents and reagents obtained from commercial sources were used without any further purification. Aqueous solutions were prepared using ultrapure water from an in-house water purification (reverse osmosis and deionization) system. Bacterial growth media and standard buffers were prepared and sterilized according to the instructions of the manufacturer without custom modifications. Bacterial growth media and cultures were handled under sterile conditions. Quantifications of protein concentrations were performed using Pierce Bicinchoninic Acid Assay (BCA) Kit (Thermo Fisher Scientific). Reversed-phase high-performance liquid chromatography (RP-HPLC) was performed with Agilent 1200 Series HPLC instruments equipped with a diode array detector. Semi-preparative and analytical C4 or C18 columns with 300Å pore sizes were purchased from Higgins Analytical. Bulk reversed-phase chromatography was performed on a Biotage system equipped with C4/C18 Sfar Bio 10g cartridges. Reversed-phase chromatography solvents are as follows: solvent A, 0.1% trifluoroacetic acid (TFA) in $H_2O$; solvent B, 0.1% TFA, and 90% acetonitrile (ACN) in $H_2O$. Mass spectra were acquired on an Agilent 1290-6545XT LC-QTOF electrospray mass spectrometry (ESI-MS) system.

**Plasmids.**

A pRK172 construct containing wild-type human alpha synuclein (α-Syn) described previously[1] was used for the expression of full-length human sequence protein. For experiments requiring the mouse α-Syn protein, custom codon-optimized gene of the mouse sequence was purchased from Integrated DNA Technologies (IDT) in a pUC19 vector. The mouse sequence was then transferred to the same pRK172 plasmid using 5' NdeI and 3' HindIII digestion sites following standard restriction digest cloning protocols. The C-terminal fragment of α-Syn (aa 91-140) was amplified from the full-length construct, then introduced into a pET42b vector using Nde I and Spe I restriction sites. The N-terminal fragment of α-Syn (1-84) was also amplified and introduced into a modified pTXB1 construct that contains a C-terminally 6xHis-tagged Ava-DnaE N137A intein[2], using NdeI and Bpu10I restriction sites and standard molecular cloning techniques.

**Expression and purification of full-length human/mouse α-Synuclein.**

BL21(DE3) competent cells (EMD Millipore) were transformed with pRK172 expression plasmids and selected over ampicillin (100 ug/mL) plates. A single colony was used to inoculate an overnight culture of Luria broth (LB) media grown at 37°C with shaking at 200 rpm. This overnight culture was expanded 1:100 into terrific broth (TB) medium and grown to an OD=0.6-0.8 at 37°C with shaking. Protein expression was induced by the addition of 0.5 mM isopropyl β-D-1-thiogalactopyranoside (IPTG) at room temperature for 16 hours. Cells were collected by centrifugation and stored at -20°C prior to lysis. Cell pellets were subjected to three rounds of freeze-thaw by submerging in liquid nitrogen for 2 minutes, followed by thawing in a 37°C incubator. The pellet was then resuspended in lysis buffer (100 mM Tris, 500 mM sodium chloride (NaCl), 10 mM beta-mercaptoethanol, 1 mM ethylene-diamine tetraacetate (EDTA), pH 8.0), and the resulting slurry was boiled at 80°C for 10 minutes. The solution was cooled to room temperature for 30 minutes, after which protease inhibitor (Roche cOmplete mini, EDTA-free) was added and allowed to incubate for another 30 minutes on ice. The mixture was clarified by centrifugation, and the pH of the supernatant was slowly adjusted to 3.5. The acidifed mixture was incubated on ice for 30 minutes and again clarified by centrifugation. The supernatant was dialyzed overnight against degassed 1% acetic acid solution at 4°C. The dialyzed proteins were then purified via RP-HPLC on a C4 semi-preparative column. Identity and purity were confirmed via analytical liquid chromatography and electrospray mass spectrometry (ESI-MS). The purified protein was lyophilized and stored as dry powder at -20°C prior to experiments.

**Expression and purification of recombinant NCL fragments.**

For the C-terminal fragment 91-140, protein expression was performed as above for full-length α-Syn but using kanamycin as a selection antibiotic for the pET42b plasmid. Lysis and purification were done via the same boiling and acid precipitation protocol described for full-length constructs. After dialysis, the N-terminal cysteine was deprotected via the addition of 100 mM methoxylamine hydrochloride, pH 3.5, at room temperature for 24 hours. The cysteine residues were then reduced by adding tris(2-carboxyethyl)phosphine (TCEP) HCl prior to final purification via C18 semi-preparative RP-HPLC. The identity of the protein fragment was confirmed by ESI-MS. The purified protein was lyophilized and stored as dry powder at -20°C prior to chemical ligation.

For the N-terminal fragment 1-84, BL21(DE3) competent cells (EMD Millipore) were transformed with the pTXB1 expression plasmid and selected over ampicillin (100 ug/mL) plates. An overnight culture from a single colony was further expanded 1:100 into fresh terrific broth media at 37°C with shaking. After reaching an optical density (OD) of 0.6-0.8, expression was induced with 0.5 mM IPTG for 16-20 hours at room temperature with shaking. Cells were harvested by centrifugation and then resuspended in a lysis buffer containing 50 mM sodium phosphate, 300 mM NaCl, 5 mM imidazole and protease inhibitors, pH 7.5. The slurry was sonicated on ice (50% amplitude, 30s on, 30s off, 3 minutes total) and clarified by centrifugation. The supernatant was applied onto pre-washed Ni-NTA agarose beads (QIAGEN) and incubated at 4°C with rocking for 1 hour. The supernatant was drained, and the beads were washed with 20 column volumes of wash buffer (50 mM sodium phosphate, 300 mM NaCl, 20 mM imidazole, pH 7.5). The intein-fusion protein was eluted with elution buffer (50 mM sodium phosphate, 300 mM NaCl, 250 mM imidazole, pH 7.5). Excess imidazole was removed by dialyzing against phosphate buffered saline at 4°C overnight. The dialyzed protein solution was clarified by centrifugation, and sodium mercaptoethanesulfonate (Mesna) was added to a final concentration of 200 mM. The pH was adjusted to pH 7, and protein thiolysis was allowed to proceed for 24-48 hours. The 1-84 thioester was purified by C4 semi-preparative RP-HPLC, and the identity was confirmed by ESI-MS. The N-terminal thioester protein was stored at -20°C as lyophilized powder prior to chemical ligation.

**Solid phase synthesis of O-GlcNAc α-Syn 85-90.**
Synthesis of the O-GlcNAcylated fragment of α-Synuclein(gS87) was performed via standard, manual fmoc-based solid-phase methods. Peptides were built on Dawson Dbz AM resin (Novabiochem). Commercially available amino acids (5 eq) were activated for 5 minutes in the presence of HBTU (4.5 eq) and diisopropylethylamine (10 eq) before coupling to the resin for 60 minutes. Following coupling and washes, N-terminal Fmoc groups were removed with 20% v/v piperidine in DMF for 15 minutes. Deprotection steps were performed twice. O-GlcNAcylated serine residues were coupled using a pentafluorophenyl (PFP) activated *per*-acetylated O-GlcNAc Fmoc-serine amino acid cassette that was synthesized and purified in-house[3]. Two equivalents of the O-GlcNAc amino acid were added to the resin-bound peptide overnight, followed by standard coupling cycles for the remaining amino acids. Residue 85 was coupled as an N-Boc-protected thioproline residue. Following completion of the peptide sequence, O-acetyl protecting groups from O-GlcNAc serine residues were removed by the addition of hydrazine hydrate (80% v/v in MeOH) twice for 45 minutes. Prior to peptide cleavage, the Dawson linker was activated with treatment of *para*-nitrophenyl chloroformate (5 equiv in $CH_2Cl_2$) for 1 hour. Cyclization was effected via incubation with excess DIEA (5 equiv in DMF) for 30 min. Peptides were then cleaved from the resin using a standard cleavage cocktail (95:2.5:2.5 TFA/$H_2O$/TIPS) for 2 hours at room temperature. Crude peptides were precipitated in cold diethyl ether, collected via centrifugation (5,000 x g, 30 min, 4°C), and lyophilized. This crude material was resuspended in thiolysis buffer (150 mM phosphate, 150 mM sodium mercaptoethanesulfonate, (Mesna) pH 6.5) and incubated at room temperature for 2 hours before purification via C18 reversed phase chromatography. Purified peptides were characterized for purity via analytical HPLC, and identity by ESI-MS.

**O-GlcNAc S87 α-Syn synthesis.**
Preparation of O-GlcNAc S87 α-Syn was performed as previously published[4]. C-terminal fragment 91-140, and O-GlcNAc S87 peptide 85-90 were incubated in ligation buffer (6M guanidine HCl, 300 mM phosphate, 25 mM TCEP, 25 mM mercaptophenylacetic acid (MPAA), pH 7) overnight at room temperature. The protected cysteine in the form of a thiazolidine at residue 85 was deprotected by the addition of 100 mM methoxylamine HCl, and adjustment of pH to 3.5. The intermediate corresponding to residues 85-140 was purified by C4 RP-HPLC and lyophilized. A second round of chemical ligation was performed by combining intermediate 85-140 and N-terminal thioester 1-140 in ligation buffer overnight. The product was purified by C4 semi-preparative RP-HPLC and lyophilized. The artificial cysteine residues were then converted to native alanines through radical desulfurization by dissolving the ligation product in degassed buffer containing 6M guanidine HCl, 300 mM NaCl, 200 mM TCEP HCl, 10% v/v t-butylthiol,

2% ethanethiol, 2 mM VA-061, pH 7. The desulfurization reaction was stirred over nitrogen at 37°C for 16 hours. The final product was purified via C4 semi-preparative RP-HPLC. Reactions and purity of intermediates and products were monitored by analytical HPLC and ESI-MS.

**Fibrillization from purified monomers.**
Lyophilized proteins were resuspended in sterile phosphate buffered saline (PBS). For Figures 4-6 and Figures S5-S9, tris buffered saline (TBS) (50 mM Tris, 150 mM NaCl, pH 7.5) buffer was used. Resuspended proteins were bath sonicated for for 15-20 minutes. Pre-formed aggregates were removed by centrifugation at 20,000 g, 4°C for 20 minutes. The supernatant was used to determine protein concentration by standard BCA measurements. The protein concentration was adjusted as indicated in the respective experiments, and the protein solutions were aliquoted into replicate experiments in 1.5 mL microcentrifuge tubes. The tubes were incubated at 37°C for 5-14 days (as indicated in specific experiments) with shaking at 1,000 rpm in an Eppendorf thermomixer.

**ThT Fluorescence.**
For the kinetic studies in Figure 2, at indicated timepoints during the aggregation, 7 µL aliquots were taken from each replicate and stored at -80°C. At the end of the aggregation period, 5 µL of each timepoint were thawed and plated onto a 96-well black, clear-bottom microplate. 195 µL of a 10 µM ThT in phosphate buffered saline solution were added onto each well. Fluorescence was measured immediately on a Biotek Synergy plate reader using 450 nm excitation, 482 nm emission wavelengths, bottom read, and gain setting 100. Fold-increase ThT values reported in the figures were calculated by dividing each measurement by the average of the time 0 baseline ThT measurements.

For routine characterization of the extent of fibrilization WT and gS87 α-Syn fibrils assembled for neuronal and animal studies, assembly solutions were verified by ThT fluorescence, as decribed previously[5,6]. The sonicated pre-formed fibrils (PFFs) were resuspended in ThT solution (50 mM glycine pH 8.5, 10 µM ThT solution), and the ThT fluorescence was measured with a FLUOstar plate reader (BMG Labtech, Germany).

**Sedimentation.**
For kinetic studies in Figure 2 and Figures S1-S2, aliquots corresponding to 10 µg of protein were centrifuged at 20,000 g for 2 hours at room temperature. The supernatants were carefully transferred onto new tubes. The same volume of 4% SDS as the original aliquot were added to the pellets, and complete resuspension of the sedimented aggregates was allowed to proceed via bath sonication and boiling for 10 minutes. 4x SDS-PAGE loading buffer (4% SDS, 40% glycerol, 0.05% bromophenol blue, 0.252 M Tris-HCl pH 6.8 and 5% β-mercaptoethanol) was added to supernatant and pellet samples, and these were boiled for an additional 10 minutes. Equal volumes of supernatant and pellet samples were loaded for each experiment. After SDS-PAGE on 12% Bis-Tris-gels and Coomassie staining, soluble and insoluble protein amounts were determined by densitometry using BioRad Image Lab software. Reported percent insoluble values were calculated as the fraction of pellet density over the sum of pellet and soluble densities.

For the analysis of monomer, oligomer, and fibril proportions (Figure S5), fibril solutions were analyzed as previously described[5]. The monomeric, oligomeric and fibrillar fractions were resuspended in 4X SDS-PAGE loading buffer 4x, and each fraction was separated on a 1 mm-thick 16% Tricine gels for 2 hours at 125 V. The proteins in the gel were stained with 0.05% of Coomassie brilliant blue diluted in 25% (v/v) isopropanol and 10% acetic acid (v/v). The gel was destained with boiling distilled water. As sonication of PFFs can lead to the release of small amounts of monomers, only PFFs preparations with residual levels of α-Syn monomers lower than 5% were used for the seeding in primary neurons[7].

**Negative stain electron microscopy.**
The ultrastructures of WT and gS87 α-Syn fibrils were analyzed by EM as previously described[5]. Activated formvar/carbon-coated 200-mesh copper grids were loaded with 3-5 µl of fibrils sample for 30 seconds and then washed three times with ultrapure water, before being negatively stained with 1% uranyl acetate

for 1-2 minutes. The excess liquid was removed, and the grids were allowed to air dry. Images were acquired on a FEI Tecnai 12 or Tecnai Spirit BioTWIN electron microscope operating at 80 kV acceleration voltage and equipped with a digital camera (FEI Eagle, FEI).

For analysis of length distribution post-sonication (Figure S5), a total of 3 to 5 fields of view for each sample were imaged, and the length of fibrils was quantified using the Image J software (U.S. National Institutes of Health, Maryland, USA; RRID:SCR_001935). Only sonicated PFFs with a 50-100 nm length were used for the seeding in primary neurons[7].

**Proteinase K cleavage.**
10 µg aliquots of protein were used for each reaction. The indicated amounts of Proteinase K were added to each sample to a total volume of 20 µL in Dulbecco's PBS (DPBS). The reactions were incubated at 37°C for 30 minutes. SDS-PAGE sample buffer was added to each reaction, and the samples were boiled for 10 minutes. Reactions were ran on 12% Bis-Tris gels using MES running buffer, and then stained with Coomassie blue.

**Seeded aggregation.**
Freshly fibrillized unmodified or O-GlcNAc S87 α-Syn were subjected to sedimentation. The pellets containing pre-formed fibers were resuspended in DPBS and the protein concentration determined by BCA. For kinetic studies, the fibril solutions were tip sonicated at 20% amplitude, 1s on, 1s off, 20 cycles total. For Proteinase K structure templating studies, fibrils were used as is.

Lyophilized unmodified human or mouse α-Syn monomers were resuspended in DPBS, bath sonicated for 20 minutes, and centrifuged at 20,000 g for 20 minutes at 4°C. The supernatant was taken and its protein concentration was determined by BCA. Monomers and PFFs were mixed at the indicated ratios, and replicate experiments were prepared in separate tubes. For kinetic experiments, reactions were agitated (1,000 rpm) at 37°C in an Eppendorf thermomixer for 24 hours. Aliquots were taken at the indicated timepoints and subjected to sedimentation assay immediately, or frozen at -80°C for later analyses. For templating studies, reactions were agitated for 7 days, then subjected to Proteinase K cleavage experiments.

***In vitro* PLK3 phosphorylation.**
Aliquots of α-Syn aggregation reactions (unmodified or O-GlcNAc S87) corresponding to 2.5 µg protein were diluted in reaction buffer (20 mM 4-(2-hydroxyethyl)-1-piperazineethanesulfonic acid (HEPES), 10 mM magnesium chloride ($MgCl_2$), 2 mM dithiothreitol (DTT), pH 7.4) then subjected to sedimentation. The supernatant was removed to a new tube, and the pellet was resuspended in an equal volume of reaction buffer, and then bath sonicated briefly. ATP (NEB, 1 mM final) and protein kinase PLK3 (Thermo PV3812, 2 ng/uL final concentration) were added to the supernatant or pellet solutions, and the reactions were allowed to proceed for 24 hours at 30°C. The reactions were quenched and solubilized by the addition of 4% sodium dodecyl sulfate (SDS) and SDS-PAGE loading buffer followed by boiling. The samples were separated via SDS-PAGE and then transferred onto nitrocellulose membrane via semi-dry transfer (Bio-Rad). The membrane was fixed with 4% paraformaldehyde (PFA) in PBS for 30 minutes and then washed 3 times with PBS. Membranes were blocked with OneBlock Western-CL (Genessee Scientific) for 1 hour at room temperature. Primary antibodies (Cell Signaling Syn204 mouse mAb or Biolegend 81a 825702 mouse mAb) were added at 1:1000 dilution and incubated at 4°C with rocking overnight. The membranes were washed in 1X tris-buffered saline with Tween (TBST) (Cell Signaling) thrice, 10 minutes each. Secondary HRP-conjugated antibodies (Jackson Immunoresearch) were added at 1:10,000 dilution and incubated at room temperature for 1 hour. The membranes were washed thrice in 1X TBST. Chemiluminescent substrate was added (Bio-Rad Clarity Western ECL), and signal captured on a Bio-Rad ChemiDoc system.

**OGA stability assay**
Bacterial OGA (BtGH84) was expressed and purified as previously described[8]. Freshly assembled fibrils or freshly resuspended monomers were diluted in PBS (25 µM final concentration) and mixed with OGA (1

uM final concentration) at a total volume of 50 µL. Reactions were incubated at 37°C for 4 or 72 hours. After the incubation time, reactions were quenched by boiling for 10 minutes. Reactions were flash frozen in liquid nitrogen and then lyophilized. For analysis, the dried protein was resuspended in 8M urea and bath sonicated for 5 minutes. Samples were injected on a C4 analytical RP-HPLC column and monitored on a gradient of 30-60% over 30 minutes. The identity of the HPLC peaks was confirmed by comparison to runs of protein standards (unmodified or O-GlcNAc S87 protein), as well as analysis by ESI-MS. Quantification was done via integration of the area under the peak using the Agilent Workstation Analysis module.

***In vitro* calpain cleavage**

Newly assembled fibrils were first subjected to sedimentation. The pellets were then resuspended in calpain reaction buffer (40 mM HEPES, 5 mM DTT, 1 mM $CaCl_2$, pH 7.2) and bath sonicated. 2U calpain (EMD 208713) were used per 2.5 ug α-Syn as titration experiments showed that this concentration gives about the same amounts of the C-terminal fragment for unmodified or O-GlcNAc S87 fibrils. Reactions were incubated at 37°C for 30 minutes, and then quenched with 2 mM EGTA. To normalize the concentrations of the truncated fibers for use in seeded aggregation experiments, SDS-PAGE and Coomassie staining were performed to enable densitometric quantitation of the truncated α-Syn band (Figure S10).

**Recombinant fibril preparation for neuronal culture experiments and animal injections**

For Figure 3 and Figure S4, after 7 days of assembly from monomers, fibrils were aliquoted and stored at -80°C. Prior to use, PFFs were diluted in PBS and sonicated for 10 cycles (1 sec on, 30 sec off, high intensity) in a bath sonicator at 10°C (BioRuptor Plus; Diagenode).

For Figures 4-5 and Figures S6-S9, after 5 days of assembly from monomers, fibrils were fragmented by sonication using a fine probe (20 sec, 20% amplitude, 1X pulse on, and 1X pulse off) directly after fibril assembly. After sonication, the PFFs were separated from the remaining monomeric solution by following the protocol published by Kumar et al., 2020[5]. The final concentration of the PFFs was quantified by the BCA protein assay according to the supplier's protocol (Pierce, Thermo Fisher, Switzerland). Sonicated α-Syn fibrils were aliquoted and stored at -80°C.

As previously established[5,7,9], α-Syn fibrils were systematically and thoroughly characterized by sedimentation, ThT binding assay, in addition to quantitative assessment of the length distribution of the fibrils before and after sonication by electron microscopy.

**Neuronal culture condition for pathology and toxicity studies.**

For experiments in Figure 3 and Figure S4, primary neuronal cultures were prepared from E16-18 CD1 mouse embryos. Dissociated neurons were plated onto poly-D-lysine coated optical bottom 96-well plates (ViewPlate; Perkin Elmer) and 60,000 cells/cm$^2$. Cultures were maintained in the Neurobasal medium supplemented with B27 (Invitrogen) that was replenished every 5 days. Sonicated PFFs diluted in sterile PBS without $Ca^{2+}$/$Mg^{2+}$ (Corning) were added at 7 days in vitro (DIV) at the indicated concentrations. Cells were fixed in 4% PFA at 19 DIV and co-labeled using antibodies against phosphorylated at serine 129 (mouse monoclonal, clone 81A, 1:10,000; CNDR), NeuN (mouse monoclonal, clone A60; Millipore) and microtubule-associated protein 2 (rabbit, 17028, 1:2,000; CNDR). Alexa Fluor-conjugated secondary antibodies mouse IgG1, IgG2a or rabbit IgG were used to visualize staining. Images were obtained using an InCell2200 scanner (GE Life Sciences).

**Intracerebral injection of PFFs.**

All housing and procedures were performed according to the National Institutes of Health Guide for the Care and Use of Experimental Animals and approved by the University of Pennsylvania Institutional Animal Care and Use Committee. The injection studies described used 2-3-month-old female B6C3F1/J mice (Stock No.100010; The Jackson Laboratories). Timed-pregnant CD1 mice for neuronal cultures were obtained from Charles River Laboratories. Animals were maintained on a 12-hour light/dark schedule and provided with food *ad libitum*.

Sonicated PFFs (5 µg total in 2.5 µL of PBS) were stereotaxically injected in the dorsal striatum using the following coordinates: AP +0.2 mm, M/L 2.0 mm, depth beneath skull 2.6 mm. Each mouse received a single unilateral injection of the indicated PFFs. At the post-injection time points indicated, mice were transcardially perfused with heparinized PBS, and brains were removed for fixation in 70% ethanol in 150 mM NaCl, pH 7.4.

**Immunohistochemistry.**

Following fixation, brains were embedded in paraffin and sectioned at 6 µm. Sections were then de-paraffinized with in xylene, followed by 1-minute washes in a descending series of ethanols (100%, 100%, 95%, 80%, 70%). Slides were then incubated in deionized water for one minute prior to antigen retrieval, as noted. After antigen retrieval, slides were incubated in 5% hydrogen peroxide in methanol to quench endogenous peroxidase activity. Slides were then incubated in blocking buffer (0.1 M Tris with 2% fetal bovine serum) and incubated with either antibody against α-Syn phosphorylated at serine 129 (mouse monoclonal, clone 81A, 1:10,000) or tyrosine hydroxylase (mouse monoclonal; clone TH-16, 1:1,000; Sigma). Primary antibody was rinsed off with 0.1 M Tris for 5 minutes, then incubated with biotinylated horse anti-mouse IgG (1:1,000; Vector BA2000) in blocking buffer for 1 hour. Sections were then washed and incubated with avidin-biotin solution (Vector PK-6100) for 1 hour. Slides were then washed and developed with ImmPACT DAB peroxidase substrate (Vector SK-4105) and counterstained briefly with hematoxylin. Slides were rinsed in water, dehydrated, and coverslipped in Cytoseal (Fisher 23-244-256). All slides were digitized using a Lamina Scanner (Perkin Elmer) in brightfield mode.

**Preparation and treatment of WT and α-Syn KO hippocampal primary neurons**

For experiments in Figures 4-5 and Figures S6-S9, WT and α-Syn KO primary hippocampal neurons were respectively prepared from WT C57BL/6JRj (Janvier, France) or α-Syn knock-out (KO) (C57BL/6J OlaHsd, Envigo, France) pups at postnatal day 0 (P0) as previously reported[10]. The dissociated neurons were plated in 6 wells plates or onto coverslips (CS) (VWR, Switzerland) or in the clear black bottom 96 well plates (Falcon) freshly coated with poly-L-lysine 0.1% w/v in water (Brunschwig). Neurons were plated at a density of 300,000 cells/mL for biochemistry analyses, 250,000 cells/mL for the ICC analyses, and 200,000 cells/mL for the HTS analyses. The α-Syn KO neurons were treated after 10 days in culture (*DIV13*) with extracellular human WT or gS87 α-Syn PFFs at a final concentration of 70 nM. The uptake, proteolytic processing, and clearance of the PFFs were followed from 14 hours after the addition of the seeds and up to 10 days. In WT hippocampal primary neurons, the WT or gS87 α-Syn PFFs seeds were added at DIV6 or DIV13 at a final concentration of 70 nM and the PFF-treated neurons were all fixed or lysed at DIV 27 as previously described[7,11–13]. PBS was used as a negative control. All procedures were approved by the Swiss Federal Veterinary Office (authorization numbers VD3496).

**Assessment of α-Syn PFF biochemistry and seeding activity in α-Syn KO and WT hippocampal primary neurons.**

PBS- or PFFs-treated α-Syn KO and WT neurons were lysed in 1% Triton X-100/ Tris-buffered saline (TBS) (50 mM Tris, 150 mM NaCl, pH 7.5) supplemented with 1/100 of protease inhibitor cocktail (Sigma-Aldrich, Switzerland), 1 mM phenylmethane sulfonyl fluoride (PMSF, Sigma-Aldrich, Switzerland) and 1/100 of phosphatase inhibitor cocktails 2 and 3 (Sigma-Aldrich, Switzerland). The cell lysates were sonicated with a fine probe (0.5 seconds pulse at 20% amplitude, 10 times) and then incubated on ice for 30 minutes and centrifuged at 100,000 g for 30 minutes at 4°C. The supernatant (soluble fraction) was collected in a new tube. The pellet was washed in 1% Triton X-100/TBS and then sonicated for (1-second pulse ON, 1-second pulse OFF, at 20% amplitude, 10 times), and centrifuged for another 30 minutes at 100,000 g. The supernatant was discarded, and the pellet (insoluble fraction) was resuspended in 2% SDS/TBS supplemented with 1/100 of protease inhibitor cocktail, 1 mM PMSF and 1/100 of phosphatase inhibitor cocktails 2 and 3. The pellet was sonicated using the fine probe (1-second pulse ON, 1-second pulse OFF at 20% amplitude, 15 times). The protein concentration in the soluble and insoluble fractions was quantified by the BCA protein assay according to the supplier's protocol (Pierce, Thermofisher). The soluble and insoluble fractions were next resuspended in Laemmli buffer 4x (4% SDS, 40% glycerol, 0.05% bromophenol blue, 0.252 M Tris-HCl pH 6.8 and 5% β-mercaptoethanol). The proteins from each fraction were separated on 1 mm-thick 16% Tricine gels (Life Technologies) for 2 hours at 125 V. The proteins were then transferred onto nitrocellulose membranes (0,2 µm, GE Healthcare) using a semidry transfer system (Life Technologies) under a constant current (25 V) and a maximum tension of 0.5 A. The membranes were then blocked for 1 hour at room temperature in Odyssey blocking buffer (Li-COR Biosciences, Germany), and probed with the relevant primary antibodies (see Table S1) overnight at 4°C. After three washes with PBS buffer containing 0.1% (V/V) Tween 20 (Fluka, Switzerland) (PBS-T), the membranes were incubated with secondary goat anti-mouse or anti-rabbit antibodies conjugated to Alexa fluor 680 or 800 dyes (Li-COR Biosciences, Germany). The membranes were then washed three times with PBS-T, and scanned on a Li-COR scanner (Li-COR Biosciences, Germany). The level of total α-Syn or pS129 α-Syn was estimated by measuring the WB band intensity using Image Studio software (Li-COR Biosciences, RRID:SCR_015795) and normalized to the relative protein levels of actin. All the experiments were independently repeated three times.

**Immunocytochemistry.**

PBS- or PFFs-treated hippocampal primary neurons were fixed in 4% formaldehyde (Sigma-Aldrich) for 20 minutes at Room Temperature and immunostained. The PFFs were detected using a total α-Syn antibody (SYN-1). The seeded-aggregates were detected using the mouse monoclonal pS129 (81a, Biolegend) or the rabbit monoclonal pS129 (MJFR-13, Abcam) antibodies and were co-stained with the ubiquitin or the p62 antibodies or with the Amytracker dye (Ebba biotech, Sweden). LAMP1 antibody was used to stain the late endosome/lysosomes. The neurons were counterstained with the microtubule-associated protein (MAP2) antibody and the nucleus with DAPI staining. All the information about the antibodies used in this study can be found in Table S1. PBS- or PFFs-treated neurons plated on coverslips were imaged with a confocal laser-scanning microscope (LSM 700, Carl Zeiss Microscopy, Germany) with a 40x objective and analyzed using the Zen software (Carl Zeiss Microscopy, Germany, RRID:SCR_013672). The PBS- or PFFs-treated neurons plated in black clear bottom 96 well plates were imaged using the IN Cell Analyzer 2200 plate reader as previously described[7,13]. For each independent experiment, three wells were acquired per condition, and in each well, nine fields of view were imaged. Each independent experiment was reproduced at least 3 times. The identification of the pS129-positive seeded aggregated formed in neurons MAP2-positive neuronal cells and the quantification of the pS129 intensity was performed using Cell profiler 3.0.0 software (RRID:SCR_007358) as previously described[7].

**Statistics.**

The data from at least 3 independent experiments were analyzed. The statistical analyses were performed using the one-way ANOVA followed by the multicomparison Tukey HSD posthoc test (compared groups are specified in the corresponding legend). The data were regarded as statistically significant at p-value<0.05.

**HSP27 fibril seeding inhibition assay**

200 µg of purified heat shock protein 27 (HSP27) C137A (prepared as previously described[14]) was denatured in 6M GnHCl, 50 mM sodium phosphate pH 8 buffer and dialysed overnight at 4 °C into DPBS for slow refolding. Protein was concentrated using 10 kDa MWCO Amicon Ultra-0.5 Centrifugal Filters and quantified by BCA. Monomeric α-synuclein was prepared by resuspending lyophilized protein in PBS buffer and bath sonicated for 15 minutes, after which the solution was clarified by centrifugation at 20,000 x g for 30 minutes. The solution was then spin filtered through Microcon DNA fast flow Ultracel regenerated cellulose columns to remove oligomers for 20min. The α-synuclein monomer and ThT stock solution were combined to a concentration of 62.5 µM each in PBS. This monomeric α-synuclein solution was divided into three conditions for treatment with different concentrations of HSP27.After incubation of the monomer-HSP mixtures at 37 °C for 30 min, seeds were added to 5% final volume for a final concentration of 50 µM monomer, and 50 µM ThT, 2.5 µM preformed fibers and respective HSP27 concentration (no HSP27, 0.125 µM HSP27, 0.5 µM HSP27).

120 µL of each solution was pipetted into the wells of a clear-bottomed black 96 well plate. Prior to assay use, the BioTek Cytation 5 instrument was preheated to 37 °C. The excitation wavelength

was set to 450nm, and emission to 482nm for ThT fluorescence monitoring over the course of 24hrs with readings taken every 5 minutes without shaking. Experiments were done in triplicate.

**Cryo-EM sample preparation and data collection.**

The cryo-EM grid preparation was performed at the Core Facilities of the Structural Biology Lab at the University of Texas Southwestern Medical Center (UTSW) using a Vitrobot Mark IV (FEI). A 3 μL aliquot of fibril sample (assembled over 14 days) was pipetted on a glow-discharged Quantifoil R 1.2/1.3, 300 mesh, Cu grid; the excess sample was removed by blotting and the grid was then plunged into liquid ethane. The cryo-EM samples were screened on either the Talos Arctica or Glacios at the Cryo-Electron Microscopy Facility (CEMF) at UTSW. The cryo-EM data was acquired using a TEM Beta, a 300 kV Titan Krios G3i electron microscope equipped with a K3 detector, at the Stanford-SLAC Cryo-EM Center (S2C2). The pixel size, frame rate, dose rate, final dose, and number of micrographs are described in detail in Table S1. The data collection was automated using the EPU 2.8 software package.

**Data preprocessing**.

All data pre-processing and processing steps were performed using RELION 3.1, unless specified otherwise[15]. The raw movie frames were gain-corrected, aligned, motion-corrected and dose-weighted using the motion correction program implemented in RELION[16]. Contrast transfer function (CTF) was estimated using CTFFIND 4.1[17]. The filaments were manually picked without discrimination using EMAN2 e2helixboxer.py[18].

**Helical reconstruction.**

The filaments were extracted using 1024-pixel boxes with a 10% inter-box distance and downscaled to 256 pixels. 2D classification was performed to select the most visually appealing classes and generate a de novo 3D initial model using the relion_helix_inimodel2d script[19]. The fibril helix was assumed to have a left-handed orientation for 3D reconstruction. Several rounds of 3D classification were carried out to determine the optimal reconstructed class. The segments from the selected class were re-extracted in 320-pixel boxes without further downscaling and underwent another round of 3D classification with an initial regularization parameter (T factor) of 4 and a large angular sampling interval of 7.5°. The T factor was gradually increased while the angular sampling was decreased stepwise with close manual monitoring of the reconstructed results. The helical twist and rise were optimized once the estimated resolution of the model exceeded 4.75 Å. After reaching a T factor of 128 and an angular sampling of 1.8, the T factor was slowly decreased to 8 with an angular sampling of 1.8 for the final map of this step. This map underwent 3D auto-refinement, then post-processed using a 10-pixel extended initial binary mask. The final resolution was estimated using the 0.143 threshold FSC between two independently refined half-maps[20].

**Model building and refinement**

The model was built in COOT using the residue sequence V77-A89 from the recombinant α-synuclein structure (pdb code 6A6B) as the template[21]. To improve the model's accuracy, multiple rounds of refinement were performed with the default settings in phenix.real_space_refine with non-crystallographic symmetry constraints, minimization_global, rigid_body, and local_grid_search[22]. The model's geometry was assessed using MolProbity, a tool integrated in Phenix. In between each refinement round, regions of the model with problematic fit or low quality were manually adjusted using COOT. This process was repeated until the model exhibited an acceptable level of stereochemistry and an adequate overall correlation coefficient with the map.

**Antibodies used in this study**

| Primary Antibody | Catalog # | Company | Clone | RRID | Host | Concentration | WB dilution | IC dilution | Epitope |
|---|---|---|---|---|---|---|---|---|---|
| anti-α-Syn total | 2647 | Cell Signaling | Syn204 | AB_2302251 | Mouse | 0.2 mg/mL | 1 :1000 | - | |
| anti-α-Syn total | 610787 | BD | SYN-1 | AB_398108 | Mouse | 0.25 mg/mL | 1:1000 | 1:1000 | 91-99 |
| anti-pS129-α-Syn | 825701 | BioLegend | p-syn /81A | AB_2564891 | Mouse | 1.0 mg/mL | 1:1000 | 1:2000 | Peptide (residues 124-134) including phosphorylated Ser129 of human α-Syn |
| anti-pS129-α-Syn (Luk lab) | - | CNDR | 81A | NA | Mouse | Not provided | - | 1 :10,000 | |
| anti-pS129-α-Syn | ab168381 | Abcam | MJF-R13 | AB_2728613 | Rabbit | 4.229 mg/mL | 1:3000 | 1:3000 | The exact sequence is proprietary |
| anti-NeuN | MAB377 | Millipore | A60 | AB_2298772 | Mouse | 1 mg/mL | - | 1 :2000 | |
| anti-TH | T2928 | Sigma | TH-16 | AB_2313844 | Mouse | 5 mg/mL | - | 1 :1000 | |
| anti-LAMP1 | ab24170 | Abcam | - | AB_775978 | Rabbit | 1.0mg/mL | - | 1:1000 | |
| anti-p62 | H00008878 | Abnova | 2C11 | AB_437085 | Mouse | 1 mg/ml | 1:1000 | 1:500 | raised against a full length recombinant SQSTM1 |
| anti-ubiquitin | Sc-8017 | Santa-Cruz | P4D1 | AB_628423 | Mouse | 0.2 mg/ml | 1:500 | 1:500 | 1-76 |
| anti-Actin | ab6276 | Abcam | AC-15 | AB_2223210 | Mouse | 2.2 mg/mL | 1:5000 | - | |
| anti-MAP2 | - | CNDR | 17028 | NA | Rabbit | Not provided | - | 1 :2000 | |
| anti-MAP2 | ab92434 | Abcam | - | AB_2138147 | Chicken | Not provided | Not tested | 1:2000 | |

| Secondary Antibody | Catalog # | Company | RRID | Concentration | WB dilution | IC dilution |
|---|---|---|---|---|---|---|
| Donkey anti-mouse peroxidase | 715-035-150 | Jackson Immunoresearch | AB_2340770 | 0.8 mg/mL | 1 :10,000 | - |
| Horse anti-mouse biotinylated | BA2000 | Vector | AB_2313581 | 1.5 mg/mL | - | 1 :1000 |
| Goat anti-mouse Alexa Fluor 680 | A21058 | Invitrogen | AB_2535724 | 2 mg/ml | 1:5000 | - |
| Goat anti-rabbit Alexa Fluor 800 | 926-32211 | Li-Cor | AB_621843 | 1 mg/ml | 1:5000 | - |
| Donkey anti-rabbit Alexa Fluor 647 | A31573 | Invitrogen | AB_2536183 | 2 mg/ml | - | 1:800 |
| Donkey anti-mouse Alexa Fluor 647 | A31571 | Invitrogen | AB_162542 | 2 mg/ml | - | 1:800 |
| Goat anti-chicken Alexa Fluor 568 | A11041 | Invitrogen | AB_2534098 | 2 mg/ml | - | 1:500 |
| Goat anti-mouse Alexa Fluor 488 | A-11029 | Invitrogen | AB_2534088 | 2 mg/ml | - | 1:800 |
| Donkey anti-chicken Alexa Fluor 488 | 703-545-155 | Jackson Immunoresearch | AB_2340375 | 1 mg/ml | - | 1:400 |
| Donkey anti-rabbit Alexa Fluor 488 | A21206 | Invitrogen | AB_2535792 | 2 mg/ml | - | 1:800 |
| Amytracker 680 | - | EBBA Biotech | Not available | 1 mg/ml | - | 1:100 |

### References


1. Waxman, E. A. & Giasson, B. A novel, high-efficiency cellular model of fibrillar alpha-synuclein inclusions and the examination of mutations that inhibit amyloid formation. *J Neurochem* **113**, 374–388 (2010).
2. Marotta, N. P. *et al.* O-GlcNAc modification blocks the aggregation and toxicity of the protein α-synuclein associated with Parkinson's disease. *Nat Chem* **7**, 913–920 (2015).
3. De Leon, C. A., Lang, G., Saavedra, M. I. & Pratt, M. R. Simple and Efficient Preparation of O- and S-GlcNAcylated Amino Acids through InBr3-Catalyzed Synthesis of β-N-Acetylglycosides from Commercially Available Reagents. *Org Lett* **20**, 5032–5035 (2018).
4. Levine, P. M. *et al.* α-Synuclein O-GlcNAcylation alters aggregation and toxicity, revealing certain residues as potential inhibitors of Parkinson's disease. *Proc Natl Acad Sci U S A* **116**, 1511–1519 (2019).
5. Kumar, S. T., Donzelli, S., Chiki, A., Syed, M. M. K. & Lashuel, H. A. A simple, versatile and robust centrifugation-based filtration protocol for the isolation and quantification of α-synuclein monomers, oligomers and fibrils: Towards improving experimental reproducibility in α-synuclein research. *J Neurochem* **153**, 103–119 (2020).
6. Fauvet, B. *et al.* α-Synuclein in Central Nervous System and from Erythrocytes, Mammalian Cells, and <em>Escherichia coli</em> Exists Predominantly as Disordered Monomer * . *Journal of Biological Chemistry* **287**, 15345–15364 (2012).
7. Mahul-Mellier, A.-L. L. *et al.* The process of Lewy body formation, rather than simply α-synuclein fibrillization, is one of the major drivers of neurodegeneration. *Proc Natl Acad Sci U S A* **117**, 4971–4982 (2020).
8. Darabedian, N., Gao, J., Chuh, K. N., Woo, C. M. & Pratt, M. R. The Metabolic Chemical Reporter 6-Azido-6-deoxy-glucose Further Reveals the Substrate Promiscuity of O-GlcNAc Transferase and Catalyzes the Discovery of Intracellular Protein Modification by O-Glucose. *J Am Chem Soc* **140**, 7092–7100 (2018).
9. Mahul-Mellier, A.-L. *et al.* Fibril growth and seeding capacity play key roles in α-synuclein-mediated apoptotic cell death. *Cell Death Differ* **22**, 2107–2122 (2015).
10. Steiner, P. *et al.* Modulation of receptor cycling by neuron-enriched endosomal protein of 21 kD . *Journal of Cell Biology* **157**, 1197–1209 (2002).
11. Volpicelli-Daley, L. A. *et al.* Exogenous α-Synuclein Fibrils Induce Lewy Body Pathology Leading to Synaptic Dysfunction and Neuron Death. *Neuron* **72**, 57–71 (2011).
12. Volpicelli-Daley, L. A., Luk, K. C. & Lee, V. M.-Y. Addition of exogenous α-synuclein preformed fibrils to primary neuronal cultures to seed recruitment of endogenous α-synuclein to Lewy body and Lewy neurite–like aggregates. *Nat Protoc* **9**, 2135–2146 (2014).
13. Mahul-Mellier, A.-L. *et al.* The making of a Lewy body: the role of α-synuclein post-fibrillization modifications in regulating the formation and the maturation of pathological inclusions. *bioRxiv* 500058 (2018) doi:10.1101/500058.
14. Balana, A. T. *et al.* O-GlcNAc modification of small heat shock proteins enhances their anti-amyloid chaperone activity. *Nat Chem* **13**, 441–450 (2021).
15. He, S. & Scheres, S. H. W. Helical reconstruction in RELION. *J Struct Biol* **198**, 163–176 (2017).
16. Zivanov, J., Nakane, T. & Scheres, S. H. W. A Bayesian approach to beam-induced motion correction in cryo-EM single-particle analysis. *urn:issn:2052-2525* **6**, 5–17 (2019).
17. Rohou, A. & Grigorieff, N. CTFFIND4: Fast and accurate defocus estimation from electron micrographs. *J Struct Biol* **192**, 216–221 (2015).
18. Bell, J. M., Chen, M., Durmaz, T., Fluty, A. C. & Ludtke, S. J. New software tools in EMAN2 inspired by EMDatabank map challenge. *J Struct Biol* **204**, 283–290 (2018).
19. Scheres, S. H. W. & IUCr. Amyloid structure determination in RELION-3.1. *urn:issn:2059-7983* **76**, 94–101 (2020).
20. Chen, S. *et al.* High-resolution noise substitution to measure overfitting and validate resolution in 3D structure determination by single particle electron cryomicroscopy. *Ultramicroscopy* **135**, 24–35 (2013).
21. Emsley, P., Lohkamp, B., Scott, W. G. & Cowtan, K. Features and development of Coot. *urn:issn:0907-4449* **66**, 486–501 (2010).
22. Adams, P. D. *et al.* PHENIX: a comprehensive Python-based system for macromolecular structure solution. *urn:issn:0907-4449* **66**, 213–221 (2010).
